# Evidence for a biological source of widespread, reproducible nighttime oxygen spikes in tropical reef ecosystems has implications for coral health

**DOI:** 10.1101/2020.03.08.982645

**Authors:** S.K. Calhoun, A.F. Haas, Y. Takeshita, M.D. Johnson, M.D. Fox, E.L.A. Kelly, B. Mueller, M.J.A. Vermeij, L.W. Kelly, C.E. Nelson, N. N. Price, T.N.F. Roach, F.L. Rohwer, J.E. Smith

## Abstract

Primary producers release oxygen as the by-product of photosynthetic light reactions during the day. However, a prevalent, globally-occurring nighttime spike in dissolved oxygen in the absence of light challenges the traditional assumption that biological oxygen production is limited to daylight hours, particularly in tropical coral reefs. Here we show: 1) the widespread nature of this phenomenon, 2) its reproducibility across tropical marine ecosystems, 3) the influence of biotic and abiotic factors on this phenomenon across numerous datasets, and 4) the observation of nighttime oxygen spikes *in vitro* from incubations of coral reef benthic organisms. The data from this study demonstrate that in addition to physical forcing, biological processes are likely responsible for increasing dissolved oxygen at night. Additionally, we demonstrate an association between these nighttime oxygen spikes and measures of both net community calcification and net community production. These results suggest that nighttime oxygen spikes are likely a biological response associated with increased respiration and are most prominent in communities dominated by calcifying organisms.

## Introduction

Between 50% and 85% of the free oxygen on Earth is produced by marine photosynthetic organisms (Tappan, 1968). While a large fraction of this production is attributed to open ocean phytoplankton, the productivity of benthic organisms often exceeds that of phytoplankton in shallow nearshore systems (MacIntyre, Geider & Miller, 1996; Daggers et al., 2018). Studies quantifying ecosystem metabolism in tropical coastal environments have historically measured net community production (NCP) using flow respirometry (Odum, 1957; Kinsey, 1985; Kraines et al., 1996) or incubation experiments (Sournia, 1976; Kraines et al., 1998). These studies have increased our general understanding of the photosynthesis-respiration equilibrium (Del Giorgio & Williams, 2007). However, they frequently lacked the temporal resolution (a few measurements per day versus one measurement every few min) and extended duration (a single day versus multiple consecutive days to months) needed to accurately identify the ecological processes driving oxygen budgets in nearshore marine ecosystems. Progress toward resolving this issue was achieved with the development of autonomous dissolved oxygen (DO) sensors (e.g., electrodes and optodes) that have produced numerous high-resolution *in situ* DO measurements in recent decades (Falter et al., 2008; Fischer & Koop-Jakobsen, 2012; Long et al., 2013; Haas et al., 2013a). Extensive DO time series data are now available for shallow, near-shore environments around the world (Viaroli & Christian, 2004; Krumme, Herbeck & Wang, 2012; Rheuban, Berg & McGlathery, 2014). The analysis and interpretation of these time series data sets is often complex because *in situ* DO concentrations depend on a multitude of known (e.g., light availability, hydrodynamics) and unknown factors (Reimers et al., 2012). Consequently, certain unexpected characteristics of such time series have remained unexplained or dismissed as equipment failure or anomalies.

One such feature of many DO time series from coral reefs and to a lesser extent other shallow, near-shore marine systems around the globe is a distinctive, pulsed increase in DO concentration at night. During such increases, or spikes, DO concentrations at night can increase to 35 µmol kg^-1^ (equivalent to 1.1 mg l^-1^) above background nighttime DO concentrations before decreasing again. In some cases, increased DO concentrations during these nightly spikes are one quarter the magnitude of daytime DO increase. Because it is generally accepted that oxygen production in the marine environment can only occur in the presence of photosynthetically active radiation (PAR) (Canfield, 2013; Lyons, Reinhard & Planavsky, 2014), these nighttime spikes were attributed to physical processes such as tidal bores, groundwater intrusion, upwelling, mixing, or thermal stratification in the water column (Kayanne et al., 2008; Takeshita et al., 2018). However, an allochthonous input of oxygen will always skew any estimate of ecosystem respiration that relies exclusively on measurements of oxygen consumption if a physical or biological origin for that oxygen is not properly accounted for (Del Giorgio & Williams, 2007). Since the DO spike at night is omitted in the classical view of biological oxygen production only being possible during the day, it is often excluded, leading to a potential underestimation of the actual biological oxygen present.

Mechanisms for biological oxygen production outside of PAR-driven photosynthesis have been proposed in recent years and serve as potential alternative explanations for nightly increases in DO concentrations, in addition to seemingly more obvious physical drivers:

### Oxygenic chlorite detoxification

by perchlorate-respiring bacteria and *oxygenic nitrite reduction* by the recently identified bacterium *Methylomirabilis oxyfera* (Ettwig et al., 2010, 2012; Schaffner et al., 2015). Schaffner and colleagues (2015) report on a chlorite dismutase enzyme cloned from the cyanobacterium *Cyanothece sp*. PCC7425 that can generate nearly 40 µM of DO s^-1^ in the presence of 1,000 µM chlorite. However, this enzyme is optimally active at pH 4.0, and minimally active to inactive above pH 7.0. The authors additionally conclude that most, if not all chlorite dismutase enzymes are optimally active at pH 6.5 or less. Ettwig et al. (2010) show that methane oxidation is coupled to oxygenic nitrite reduction in the bacterium *M. oxyfera*, isolated from anaerobic sediments. Their experimental setup demonstrated up to 70 nmol of DO generated over 6-8 hours by *M. oxyfera* from a substrate containing 2 mM nitrite. However, the substrate used to produce this quantity of oxygen also contained propylene, as little measurable oxygen was generated from a methane substrate (the bacteria’s preferred carbon source which is metabolized via consumption of generated oxygen). Overall, for oxygenic processes involving either nitrite or chlorite to be responsible for nighttime DO spikes on coral reefs, either compound (or [per]chlorate) would need to be present in millimolar quantities, which has to date not been shown.

### Far-red light photosynthesis

(Gan & Bryant, 2015). Far-red light photosynthesis in cyanobacteria has been proposed over the last decade as an adaptation to environments enriched in light with wavelengths greater than 700 nm, well outside the normal PAR spectrum. Such environments include benthic surfaces shaded by other photosynthesizers, interior layers of stromatolites and biofilms, karst caves, and sediments (Averina et al., 2018). Gan et al. (2014) used a systems biology approach to describe the cyanobacterium *Leptolyngbya* sp. strain JSC-1 and its adaptation to far-red light by synthesizing two different types of chlorophyll: Chl *d* and Chl *f*. Not only did this cyanobacterium remodel its photosynthetic pathway under far-red light, it grew well, leading the authors to conclude that it was both producing oxygen and fixing carbon as if it were exposed to light in the normal PAR spectrum. However, far-red light has very poor penetration in the water column, limited to 10 m depths or less, and is predominantly available during the day (Gan & Bryant, 2015).

### Reactive oxygen species (ROS) detoxification

ROS can regenerate molecular oxygen from oxygen radicals and hydrogen peroxide produced during aerobic respiration (Guzy & Schumacker, 2006). Corals have recently been documented to release up to 1.8 µM of hydrogen peroxide (H_2_O_2_) over a period of 20 min when exposed to physio-chemical stimuli such as occurs during filter feeding (Armoza-Zvuloni et al., 2016). The half-life of H_2_O_2_ in marine environments is estimated to be on the order of hours to days, with up to 80% of its breakdown occurring via microbial-derived catalase (Zinser, 2018). Assuming these parameters, an estimated 0.7 µM of oxygen could potentially be produced via H_2_O_2_ diffusion into and DO diffusion out of microbial cells over the course of a night (assuming the catalase reaction stoichiometry of 2 H_2_O_2_ → 2 H_2_O + 1 O_2_).

Several possibilities thus exist for biological oxygen production in the absence of PAR. In this study, we describe nighttime DO spikes occurring on coral reefs around the world and explore potential mechanisms for their biological origin. We reviewed the literature and conducted *in situ* DO time series, mesocosm experiments *in situ*, and controlled laboratory experiments *in vitro*. Several statistical modeling approaches were used to elucidate drivers of DO spike occurrence and describe features of the DO spikes themselves, namely the height of the spikes and times of occurrence. Data collected concurrently with DO measurements (e.g., pH, temperature, and water current profiles) and more infrequent measurements (e.g., total alkalinity, dissolved inorganic carbon, and offshore hydrodynamic and weather parameters) were combined in these models, to determine if nighttime DO spikes are widespread, frequent, and explainable by single or combinations of physical and environmental variables. We found that a limited set of closely-linked, biologically-mediated variables can accurately describe nighttime spikes in DO. Furthermore, nighttime spikes in DO were observed under controlled laboratory conditions, ruling out many physical forcings (e.g., tidal flux… etc.). These findings are then discussed in the context of ongoing threats to coral reef health and the overall implications of biological oxygen release at night.

## Methods

### Review of Literature

We performed an extensive literature search to determine the prevalence of nighttime spikes in DO from published *in situ* time series datasets. In total, >3000 papers were reviewed using ISI Web of Science, Google Scholar, and the University of California San Diego Research Data Collections (http://library.ucsd.edu/dc/rdcp/collections) with the keywords ‘coral oxygen’, ‘marine nighttime oxygen’, ‘marine oxygen’, ‘aquatic oxygen’, and ‘marine calcification’ with publication dates between 1970 and 2015. Visually screening published figures was found to be the most efficient high-throughput method of determining whether or not a publication contained DO time series data that spanned both day and night times with high enough resolution to observe a nighttime DO spike. Up to the first 1,000 publication hits for each keyword phrase were screened for figures of *in situ* DO time series plots with three or more consecutive time-point measurements per night spanning at least 24 hours. After identifying plots that met our criteria, we distinguished nighttime DO spikes in those plots as DO concentration increases in the absence of PAR, lasting at least one hour before decreasing again. In total, 28 (20 temperate, 8 tropical) relevant datasets were identified, where 19 showed an increase and subsequent decrease in DO at night generally lasting 4-6 hours (Table S1).

### Datasets Used for Statistical Analyses

Our *in situ* datasets represent autonomous multi-sensor sonde (hereafter referred to as ‘sensor’) deployments across the central Pacific and Caribbean basins spanning the years 2010 - 2015. DO datasets were obtained from enclosures deployed during September 2011 on the island of Mo’orea, French Polynesia using a single sensor inside collapsible benthic isolation tents (cBITs - Figure S1, after Haas et al. (2013)). A similar method was used in 2010 and 2013 in the Line Islands (11 Pacific islands stretching 2,350 km northwest-southeast across the equator), hereafter referred to as the Line Islands cBIT data set. *In situ* DO and temperature surveys (i.e., sensor deployments where no enclosure was used) were taken in September, 2014 at Palmyra Atoll in the northern Line Islands (Takeshita et al., 2016), hereafter referred to as the Palmyra BEAMS (Benthic Ecosystem and Acidification Measurement System) dataset, and in May, 2015 on the island of Curaçao in the southern Caribbean. Additional measurements of percent benthic cover, total alkalinity (TA), and dissolved inorganic carbon (DIC) were taken for the Line Islands cBIT and Palmyra BEAMS data sets. The saturation state of aragonite (Ω aragonite) was calculated from daily TA and DIC samples for each data set, as described under the heading for each.

#### Line Islands cBITs 2010 and 2013

The Northern Line Islands consist of 5 individual islands spanning latitudes from 6°24’N to 1°53’N in a northwest to southeast trend. Atoll and fringing reef structures dominate the marine terrain around each, consistent with the whole of the Line Islands chain. Research was conducted across these islands from October 24 to November 23, 2010 using the cBIT setup previously described (Figure S1), with a mean (± standard deviation [SD]) daily PAR measurement of 328 ± 175 µmol photons m^-1^ s^-1^ and water temperature of 26.5 ± 1.3 °C collected at 5 min intervals and 10 m depth using a LICOR (LI-COR, Inc., www.licor.com) and a MANTA multiprobe sonde (configured the same as the Mo’orea cBIT deployments), respectively. The Southern Line Islands are an additional 6 islands of the Line Islands chain spanning 0°22’S to 11°26’S, making the whole of the Line Islands one of the longest island chains in the world (2,350 km from north to south). Research studies were conducted across these islands from October 18 to November 6, 2013, with a mean ± SD daily PAR measurement of 312 ± 214 µmol photons m^-1^ s^-1^ and water temperature of 28.1 ± 0.5 °C (collected as described for the Northern Line Islands). cBITs, each containing one multi-probe sonde, were deployed at 10 m on the fore-reef habitat in all the Line Islands (6 cBITs per island, across 11 islands). Percent benthic cover was estimated from photoquadrats, and the percentages for various organisms classified as either ‘Calcifiers’ (calcifying algae such as *Halimeda* sp., crustose coralline algae, and hard corals), or ‘Non-Calcifiers’ (fleshy macroalgae, turf algae, soft corals, and corallimorphs). TA and DIC samples were collected on 24-hour intervals starting midday following the procedure outlined by Haas et al. (2013), where a pump was placed inside the cBIT and a line fed out of the cBIT underneath the skirt. Water samples were placed in 300 ml borosilicate glass containers, poisoned with mercuric chloride, and sealed with glass stoppers. TA and DIC were analyzed using standard procedures (Dickson et al., 2007). pH on the total scale and Ω aragonite were calculated using CO2SYS with equilibrium constants from Lueker, Dickson & Keeling (2000).

#### Palmyra BEAMS 2014

A single island research study was carried out from September 8 to 24, 2014 on Palmyra Atoll, the second northernmost island of the Line Islands (5°52’N 162°6’W) as previously described (Takeshita et al., 2016). Briefly, a benthic flux method was used to determine the vertical gradients of DO and pH starting at the benthos. These data was then used to calculate time series fluxes of net community production (NCP, using the gradient of DO), and net community calcification (NCC, using the gradient of pH). Water current speed and direction measurements were also made alongside the vertical gradient measurements, and subsequently used in the calculation of NCP and NCC. A time series of Ω aragonite was also calculated from the gradient of pH and verified by daily TA and DIC measurements as described by Takeshita et al. (2016). Percent benthic cover was estimated from photoquadrats, and the percentages for various organisms binned into ‘Calcifiers’ or ‘Non-Calcifiers’ as described for the Line Islands cBIT data. These additional variables make this data set the most comprehensive data presented here in terms of site level physical and biological variables.

#### Mo’orea cBIT

Sites at the island of Mo’orea (17°48’S 149°84’W) were monitored from September 1 through 22, 2011. cBITs were deployed for 36 hours at 5 m depth on the back reef habitat of Mo’orea for the purpose of isolating the benthic water column from the surrounding seawater, after the methods described by (Haas et al., 2013b). Briefly, each cBIT contained a MANTA multiprobe sonde (Eureka Water Probes, www.waterprobes.com) with sensors measuring and logging pH, redox potential (ORP), conductivity, dissolved oxygen (DO), and temperature on 5 min intervals.

#### Curaçao

The island of Curaçao (12°7’N 68°56’W) is located approximately 64 km northeast of the Venezuelan coast on the southernmost edge of the Caribbean tectonic plate. It is a semi-arid island surrounded by fringing reefs, with greater coral diversity and coral coverage than much of the Caribbean. Research on this island was conducted out of the CARMABI Research Station from April 14 to May 28, 2015. A mean ± SD temperature of 26.7 ± 0.1 °C was collected as described for the previous islands. A single MANTA multiprobe sonde was deployed in Curaçao at 10 m of depth approximately 200 m offshore of a desalinization plant located at 12°6’N, 68°57’W. The MANTA was set up to autonomously log parameters per the deployment for Mo’orea and the Line Islands, with the exception that no cBITs were used. No PAR data was taken due to the lack of an appropriate PAR sensor during this expedition.

### Physical and meteorological data

Oceanographic and meteorological data were used to assess the contribution of global scale physical processes. NOAA weather buoy data (http://www.ndbc.noaa.gov) from buoys stationed at 155°W and 8°N-8°S over the dates listed for Northern and Southern Line Islands cBIT deployments was used to obtain pressure at 300 and 500 m of seawater depth (indicative of potential offshore currents/upwelling), as well as wind speed and direction. Moon phase and intensity data were obtained from naval astronomical charts (http://aa.usno.navy.mil/data/docs/MoonPhase.php) and used as a proxy for both tidal forcing and moonlight.

### Statistical Analysis of Datasets

All time series data were analyzed using MatLab (R2018a, MathWorks, Inc. https://www.mathworks.com) and R (v3.5.1, R Core Team, https://www.R-project.org/). DO concentrations in μmol kg^-1^ seawater mass (the oceanographic standard unit for DO) were calculated from percent saturation, temperature and conductivity measurements using functions from SEAWATER Library v. 3.3 (http://www.cmar.csiro.au/datacentre/ext_docs/seawater.htm) and Gibbs Seawater Oceanographic Toolbox (http://www.teos-10.org/pubs/gsw/html/gsw_contents.html). For reference, DO values between 200 and 210 μmol kg^-1^ are equivalent to 100% air saturation at temperatures normally observed in tropical seawater (from 28 to 24 °C, respectively). Data collected via MANTA sondes were normalized to the overall average of the first 30 min of all six sensors in the Line Islands deployments (using cBITs). Additionally, daily pH values on the total H^+^ ion scale were used to calibrate the pH time series by baseline regression between each discreet pH value.

Data in each time series data set were collected at different time intervals and at the site (at the location of the sensor/sampling) or island level (one set of data points for an entire island or many islands) (Table S2). In order to analyze all data together, time points were linearly interpolated onto the most frequent time scale for hourly and daily time points.

### Nighttime DO Spike Identification

An algorithm for defining a nighttime DO spike was empirically derived based on DO spike parameters observed in the literature and the data sets presented in this study. Time series data were separated into night and day times using either PAR values or sunset and sunrise times, and then smoothed to reduce noise using a moving average filter with a sliding window of 2 hours (Figure S2A). Next all nighttime DO spikes with a prominence (height) greater than or equal to 0.5 μmol kg^-1^ and a duration (defined as the width of the spike at half its height, Figure S2B) of at least 1 hour were identified computationally using MatLab function findpeaks. A value of 0.5 μmol kg^-1^ was selected as the smallest possible spike height that was at least double the maximum sensitivity threshold of the DO sensors used (sensitivity threshold of 0.2 μmol kg^-1^ for MANTA multi-probe sondes used in all data sets except Palmyra 2014; 0.1 μmol kg^-1^ as reported by Takeshita *et al*. for the Palmyra 2014 data). Data analyzed using this algorithm were manually checked to determine the validity of any identified nighttime DO spikes. Varying height and duration values, as well as the size of the time series smoothing window did not change the algorithm’s ability to reliably detect a spike in DO concentration.

### Data Patterns at the Time of a DO Spike

To test potential mechanisms underlying nighttime DO spikes, we utilized a suite of data collected *in situ* concurrently with DO. The simultaneous change of various oceanographic parameters such as temperature, salinity, current direction and current speed could indicate whether or not an DO spike coincides with changes in the overlying water mass (Kayanne et al., 2008). Variables such as benthic cover, DO and PAR data from the previous day, and pH changes associated with biological activity (such as calcification) could also lead to a biological source of nighttime DO spikes. Therefore, we analyzed our own datasets to assess whether the presence or absence of nighttime DO spikes could be classified using measurements of multiple variables at the exact time of a spike. Calculating the first (Δ) and second (ΔΔ) derivatives of these variables allowed the degree of change occurring, the time of local maxima or minima, and the time of inflection points to be taken into account as well (Figure S3). Additional discrete values for benthic cover, as well as DO and PAR data from the previous day were included.

#### Random forests

Random forests analysis utilizes multiple tree-clustering algorithms to select the most important parameters when predicting values for numerical regression or classification based on non-numerical data. One major benefit of this method is that it does not require any *a priori* information about the distribution or frequency of values in a dataset, unlike many linear modeling approaches. When carried out with multiple permutations of the data, a probability for how reliable the results are can also be obtained. This technique was employed to discover which, if any variables might be able to best predict the presence or absence of a DO spike. After identifying the most prominent DO spike per night across all datasets, all data values at the times of these spikes were used as potential classification predictors for DO spike occurrence. The frequency of nighttime DO spikes for a particular time series set (e.g., all time series data for one site) was binned into 1-hour time points across a 12-hour time span (sunset to sunrise). A time point that represents the middle of the most frequent time bin was used as a proxy for the time of a DO spike on nights where no spike was identified. Values at these time points were used as predictors for the absence of a DO spike. The first and second derivatives of time series variables, as well as nightly sums of variables (total amount measured per night as determined by numerical integration of the area under the curve), and values from the previous day (mean DO, mean PAR, and the ratio of integrated daily DO to integrated daily PAR) were also included. Percentages of benthic cover estimated from photoquadrats were included as discreet values for each site.

Combining datasets increased the power of analyses while testing potential drivers across as many examples of nighttime DO spikes as possible. The Line Islands cBITs and Palmyra BEAMS datasets were combined due to the large number of variables each have in common (Table S2). However, no percent benthic cover, Ω aragonite, or NOAA buoy data were available for the Mo’orea or Curaçao datasets, limiting any assessment of potential drivers to pH, temperature, and DO. Therefore, these two datasets were screened for nighttime DO spikes, but ultimately not used for further analyses.

Random forests classification analysis was performed using package rfPermute in R (Archer, 2019). Presence/absence predictor data were analyzed using 3000 permutations, and significant p-values (< 0.05) for all predictors calculated. Overall strength of the classification analysis was assessed using the out-of-bag error rate (i.e., the rate of misclassification). Classification was performed at the site level and site-plus-island level in order to reduce any bias caused by uniformity of island level variables across individual sites. The top predictors in terms of mean decrease in accuracy of classification (i.e., if such predictor was excluded) and significance were then selected and used as the only predictors for a second round of random forests classification. Those predictors that retained a high value of the mean decrease in accuracy score were then considered top predictors.

#### Structural equation models

An advantage of the random forests analysis as employed here is the ability to select potentially important variables *a priori* from a large data set for further analysis. Using the top predictors of DO spike occurrence, more detailed models were constructed that combined multiple variables into nested structures to analyze interactions between variables in what is known as a structural equation model (SEM) (Lefcheck, 2016). These top predictors were subsequently used as both predictors and response variables in a set of linear mixed effects models for the same presence/absence data. Original data (not the first and second derivatives or any ratios) were transformed via hyperbolic arcsine and derivatives and ratios were scaled by one or two orders of magnitude to provide a better fit for linear modeling. Correlations between pairs of variables per site were penalized using a spherical correlation matrix based on latitude, longitude and individual site designation (R package nlme, function corSphere). Island name was used as a random effect for all models. A set of nested models were constructed, starting with presence/absence as the response for a generalized linear mixed-effects model using a binomial distribution (R package lme4, function glmer). The top predictors from random forests classification were used as the predictors in this model. Each top predictor was also used as the response variable in a linear mixed-effects model with a Gaussian distribution (R package lme4, function lme). Q-Q plots of the residuals in each model were used to confirm a Gaussian distribution. The predictors for these models were selected from the remaining variables in the data. Each model was then combined into a structural equation model (R package piecewiseSEM) to assess the influence each predictor has on each response variable. Predictors were added or removed from models based on their significance and coefficient strength using the missing paths predicted from the structural equation model. Models were optimized using this missing paths strategy to achieve the highest possible conditional r-squared value (the ratio of variance in the data explained by both the fixed and random effects in the model) and the lowest possible Aikake information criteria score (AIC) (Shipley, 2013). A similar process was carried out on only the data with a DO spike present, using height and time as response variables in two separate starting models instead of presence/absence.

#### Robust linear regression

Robust linear regressions were performed with pairs of variables selected from the SEM analyses with estimation using Tukey’s biweight (R package MASS, function rlm) (Venables & Ripley, 2002) and corresponding bootstrapped 90th percentile and 95th percentile confidence intervals (90% and 95% CIs) for the slope using 1,000 bootstrap replications. Confidence intervals (CIs) for the robust regression analyses only describe the confidence in the slope of the regression line. If the CIs at 90% and 95% include zero, this means that a basic description of the slope as either negative or positive cannot be made. The robust regression itself can still draw a best fit line indicating a linear relationship, but the reliability of said line’s slope cannot be ascertained without bootstrapped CIs. Regression lines are drawn as solid lines if the CIs at either 90% or 95% do not include zero, while a dashed line indicates the inclusion of zero in both CIs.

### Laboratory Incubations Using Wild and Cultured Reef Organisms

To determine if nighttime DO spikes observed *in situ* could be isolated in the laboratory, incubations were carried out using both wild collected and aquarium-cultured reef organisms. Wild collected organisms were obtained from the reefs of Curaçao during two separate sets of experiments, the first over April-May, 2015 and the second April-May, 2016. Aquarium-cultured Coral fragments reared for aquaculture purposes were grown in a 1,000 gal recirculating artificial reef system at San Diego State University, San Diego, USA for 6 months and used for incubation experiments over Feburary-March, 2018.

Transparent polycarbonate tube design, with rubber gasket sealed lids on either end were used as incubation tanks. Each incubation tank measured 7.5 cm in diameter by up to 50 cm tall and held volumes of 2, 1.5 and 1 l of seawater. Different incubation tank volumes were used for different amounts of organism biomass (estimated from seawater displacement volume) and to accommodate different sensor and water bath sizes. Tanks were installed in a temperature-controlled water bath that was placed in a temperature-controlled room (mean ± SD of 24.0 ± 0.5 °C for the water bath used in Curaçao, and 25.0 ± 0.5 °C for water baths used at SDSU). No external or internal water flow was allowed in any incubation tank during the incubation period to minimize the possibility of introducing external oxygen. For dark incubations, light intensity measured inside the water bath by HOBO data loggers showed 0.0 lux, an effectively lightless environment. Diurnal light cycle incubations were carried out in a fully dark environment with all light coming from a combination of LED and T4 fluorescent lights on a 12-hour cycle (Figure S4).

Multiple sensor types were deployed to measure DO and temperature during the incubations. These included MANTA multisensor sondes (sensitivity thresholds of 0.2 μmol DO kg^-1^and 0.01 °C), a single-channel fiber-optic oxygen sensor ([sensitivity threshold of 0.25 μmol DO kg^-1^] PreSens Precision Sensing GmbH, www.presens.de) combined with a HOBO temperature logger ([sensitivity threshold of 0.14 °C] HOBO Pendant Temperature/Light 8K Data Logger, Onset Computer Corp., www.onset.com), and a handheld optical sensor set to continuously log measurements ([sensitivity thresholds of 0.2 μmol DO kg^-1^and 0.1 °C] HACH HDQ Portable Meter with optical oxygen sensor, HACH Company, www.hach.com).

Several different benthic components were collected from the reefs of Curaçao and brought into the lab during April-May, 2015. Benthic components included: fine grained sand/sediment (‘Sediment’), dead coral rubble covered in turf algae (‘Turf’), bare dead coral rubble (‘Rubble’), crustose coralline algae (‘CCA’), and a mix of the aforementioned benthic components (25% of each) (‘Mixed’) (See Table S4 for a list of all samples incubated). Sample composition was determined by visual assessment of the surface area of the sample, where the sample was classified as one of the five types per the dominant (>75% of surface area) sample type present. Benthic samples comprising the five basic types listed were collected from varying depths at several locations across the island of Curaçao (Table S9 & S11). All samples were collected in polycarbonate incubation tubes filled with natural seawater, sealed and transported to CARMABI research station where they were immediately transferred to new incubation tubes filled with seawater from the flow-through system and placed in the incubation chamber, except for certain CCA samples. CCA samples collected deeper than the intertidal zone were allowed to recover for 48 hours in a low-light aquarium with flow before experiment began in order to minimize the stress of being chiseled off the reef during collection. No other samples were collected by chiseling.

Differences in volumetric displacement (a rough proxy for biomass) were used for each benthic type to determine if the nighttime DO spikes occurred due to a specific volume of sample material. Samples were either incubated in singlet, duplicate or triplicate depending on the amount of sample available. A total of four control samples were incubated: a tap-water control to check the abiotic DO to temperature correlation, a water column control collected from the reef at 10 m depth, and a water column control from the surface water to observe the DO variability in the water column alone. Furthermore, dry rubble exposed to direct sunlight for 12 hours then submerged in seawater was used to determine whether DO changes resulted from rubble removed from a typical reef system. Tubes were oriented vertically with a sealed bottom lid and open top for access by the sensors. Additionally, the surface area of the seawater inside each tank exposed to air was minimized as best as possible by the position of the sensors at the top and vertical orientation of each tube, blocking direct seawater-air contact.

A similar incubation setup was used during the incubation experiments carried out over April-May, 2016. Incubation chambers were similar as before but could be fully sealed using customized lids that allowed either a fiber-optic optode (PreSens) or a multisensor sonde (MANTA) to be placed in a fitted port in the lid. This ensured the creation of an air tight seal, verified by incubating deoxygenated water for 12 hours. Total chamber volumes were 1 l for the chambers using a PreSens optode and 1.5 l for those using a MANTA array. This was done to help compensate for water displaced by the relatively large MANTA array compared to that of the thin fiber-optic optodes. The incubation chambers were placed vertically in a water bath covered with a tent of light-blocking fabric over a PVC frame. Aquarium lights (maximum PAR of 300 µmol photons m^-2^ s^-1^) on a digital timer were attached to the inside of the PVC frame, allowing for simulation of a controlled diurnal cycle that mimicked the sunrise and set times for the area.

Incubations of aquarium-cultured coral were carried out over February-March, 2018 using exclusively MANTA sensors in 1 l polycarbonate incubation tubes. This was done so that the entire tube and attached sensor could be placed horizontally and submerged underwater. Fragments of the hard coral of *Montipora capricornis* were incubated either in the dark as described for the incubations in Curaçao, or in the aquarium where they were originally growing such that they maintained the same temperature and light cycle.

All incubation DO time series data were analyzed using the aforementioned computational algorithm to identify any DO spikes. Additionally, for any open-top incubations, Fick’s first law of diffusion was used to calculate the amount of oxygen potentially introduced via diffusion across the air-water interface (Wanninkhof et al., 2009) and that amount was subtracted from the DO measurements. See Supplemental Methods for a detailed description of these calculations.

## Results

### Review of Literature

An extensive literature search identified 28 (20 temperate, 8 tropical) studies that have published figures of DO time series spanning 24 hours or more between 1970 and 2015. Including an additional 13 datasets collected during this study, 78% (32 out of 41) document an increase and subsequent decrease in DO concentration during the night lasting between 4 and 8 hours, with 63% (20 out of 32) of those studies originating in the tropics (Figure 1, Table S1). Quantitative comparisons across these datasets are difficult due to differences in methodologies and reported DO units (e.g., oxygen flux rates, isotopic ratios, percent air saturation, mmol m^-2^ d^-1^), and calculating DO values for each using a consistent metric would require access to numerous unpublished datasets. However, qualitative details can be observed in the published figures and data. Two studies on the Island of Mo’orea (Sournia, 1976; Campion-Alsumard et al., 1993) (Figure 1, inset numbers 29-30) identified a nighttime DO spike, but did not discuss it. Hydrodynamics were suggested to underlie nighttime variability in DO off the coast of Japan, bringing more oxygenated offshore water onto the reef (Kayanne et al., 2008) (Figure 1, number 12 inset). Evidence in support of this hypothesis includes shifts in pH, temperature, and tides that occurred simultaneously with the DO spike, indicative of changing hydrodynamic conditions at the study location.

**Figure 1.**
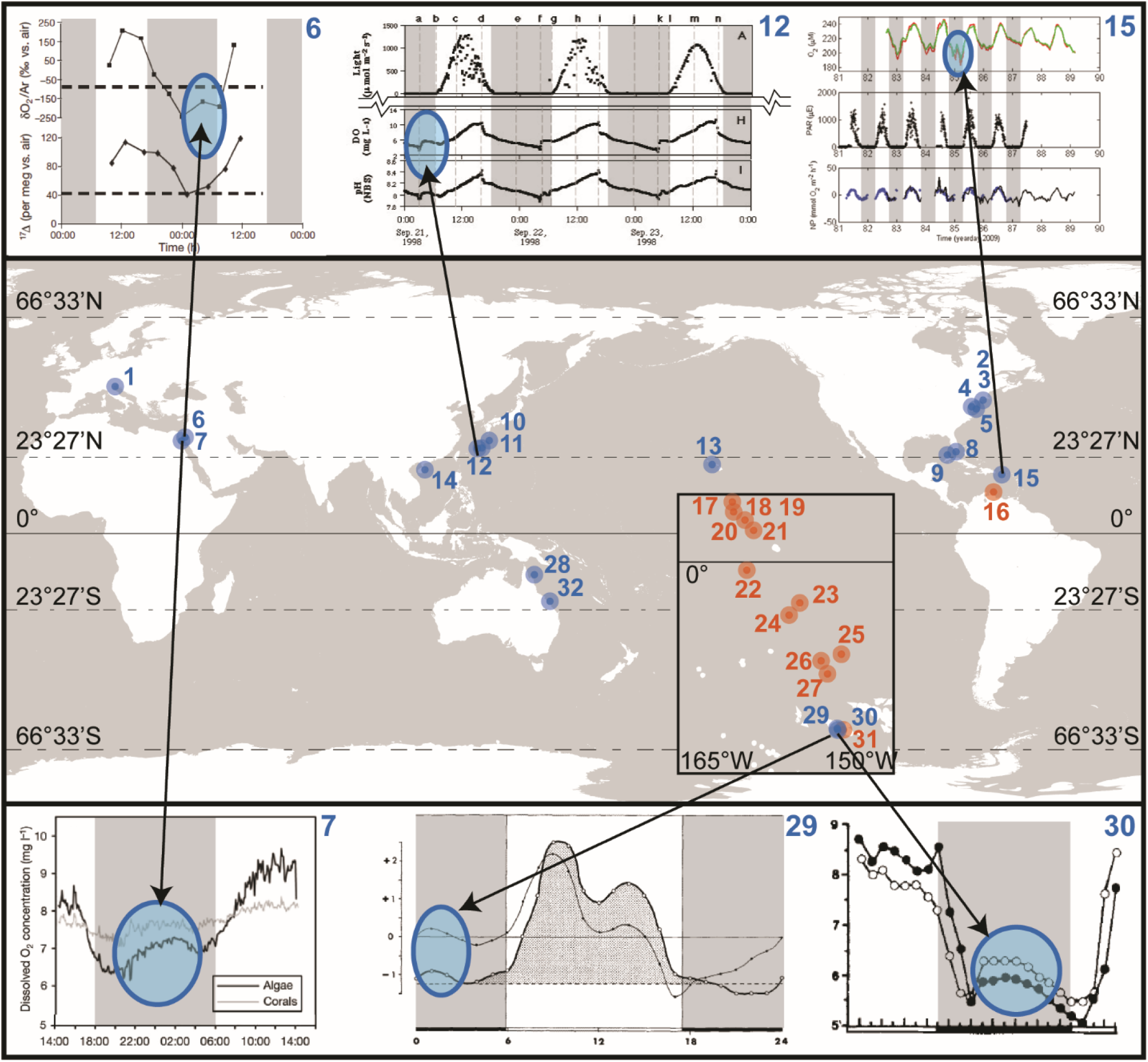
Nighttime spikes in dissolved oxygen concentrations are global phenomena. Each number corresponds to a dataset identified during a literature search (blue dots and numbers) or a dataset presented in this study (orange dots and numbers) consisting of dissolved oxygen concentration measurements across day and night times for at least 24 hours. The equator, tropics, and polar latitudes are labeled.

Methods can indicate whether the DO increase results from physical or biological processes. Because argon’s dissolution properties in seawater are very similar to those of oxygen, changes in oxygen:argon ratios (δO_2_:Ar) can be used to distinguish between biologically produced oxygen increases and atmospheric dissolution of oxygen. Luz and Barkan (2009) used this method to calculate net oxygen production in the Red Sea (Figure 1, number 7 inset) and showed a spike in δO_2_:Ar between 01:00 a.m. and 05:00 a.m., which supports biologically derived net oxygen production at night. The authors do not discuss this observation or any implications related to it, however. These studies illustrate that the origin of DO spikes can be attributed to both biological and physical processes, but also that they are often ignored. Nevertheless, the observation of a spike in DO at night in numerous published datasets appears to be a common phenomenon across the tropics, though the underlying mechanisms have never been specifically investigated.

### Repeatability of Nighttime Oxygen Production *In Situ*

We identified nighttime DO spikes in all our datasets with varying degrees of frequency, height, duration, and time of occurrence (Table S3 & S4). The percentage of nights with a DO spike compared to all nights observed is 81% (231 out of 284, Table S3), just above the 78% observed in the literature. However, the proportion of nights during which DO spikes were observed differed among locations: Curac□ao – 67% (4 out of 6); Palmyra BEAMS – 71% (44 out of 62); Mo’orea – 76% (41 out of 54); Line Islands cBITs – 88% (142 out 162).

The highest DO spike across all our datasets was measured at 35.1 µmol kg^-1^ on the island of Mo’orea (Table S4, Figure S5B – orange asterisk). DO spikes on Mo’orea’s reefs were on average the highest and longest lasting of any dataset, but also the most variable (mean ± SD of 8.6 ± 7.4 µmol kg^-1^ and 3.0 ± 1.3 h). Height for DO spikes in the remaining datasets ranged from a mean (± SD) of 1.5 ± 1.0 to 7.3 ± 7.1 µmol kg^-1^, and the mean spike width for all sites was around 2.6 ± 1.2 h. The widest DO spike occurred on Vostok island in the Southern Line Islands chain (8.0 h on Oct. 24, 2013 02:30). The smallest spikes in terms of both height and width were observed on the island of Curac□ao (mean ± SD of 1.5 ± 1.0 µmol kg^-1^ and 1.9 ± 0.4 h, and Figure S5A). Median values for both height and width are always smaller than the mean, indicating the data skew toward smaller values for overall DO spikes. Diel fluctuations in DO concentration also varied greatly from site to site, with no obvious upper or lower threshold limiting the occurrence of a nighttime DO spike. Average DO concentration reached during the spikes varied from 137.0 ± 20.0 µmol kg^-1^ at Mo’orea to 188.0 ± 11.4 µmol kg^-1^ in the Line Islands. In sum, at every location we studied, nighttime DO spikes we documented occurring across multiple nights, with each set of DO spikes exhibiting location-specific local characteristics.

The Line Islands cBITs and Palmyra BEAMS datasets combined represent 79% of all observations across all datasets (224 out of 284, Table S3), with 83% of those being positive identifications of a DO spike (186 out of 224). Taken in conjunction with the fact that these two datasets contain the most shared variables across all of the datasets (Table S2), the subsequent analyses focus exclusively on these data. The number of nights with and without DO spikes shows a latitudinal trend where most of the nights without DO spikes occur nearer the equator for the Line Islands cBITs (Fanning to Millennium, Figure 2A). Most spike heights are between 0.5 and 10 µmol kg^-1^ and occur more than 4 hours after sunset (Figure 2B & 2C).

**Figure 2.**
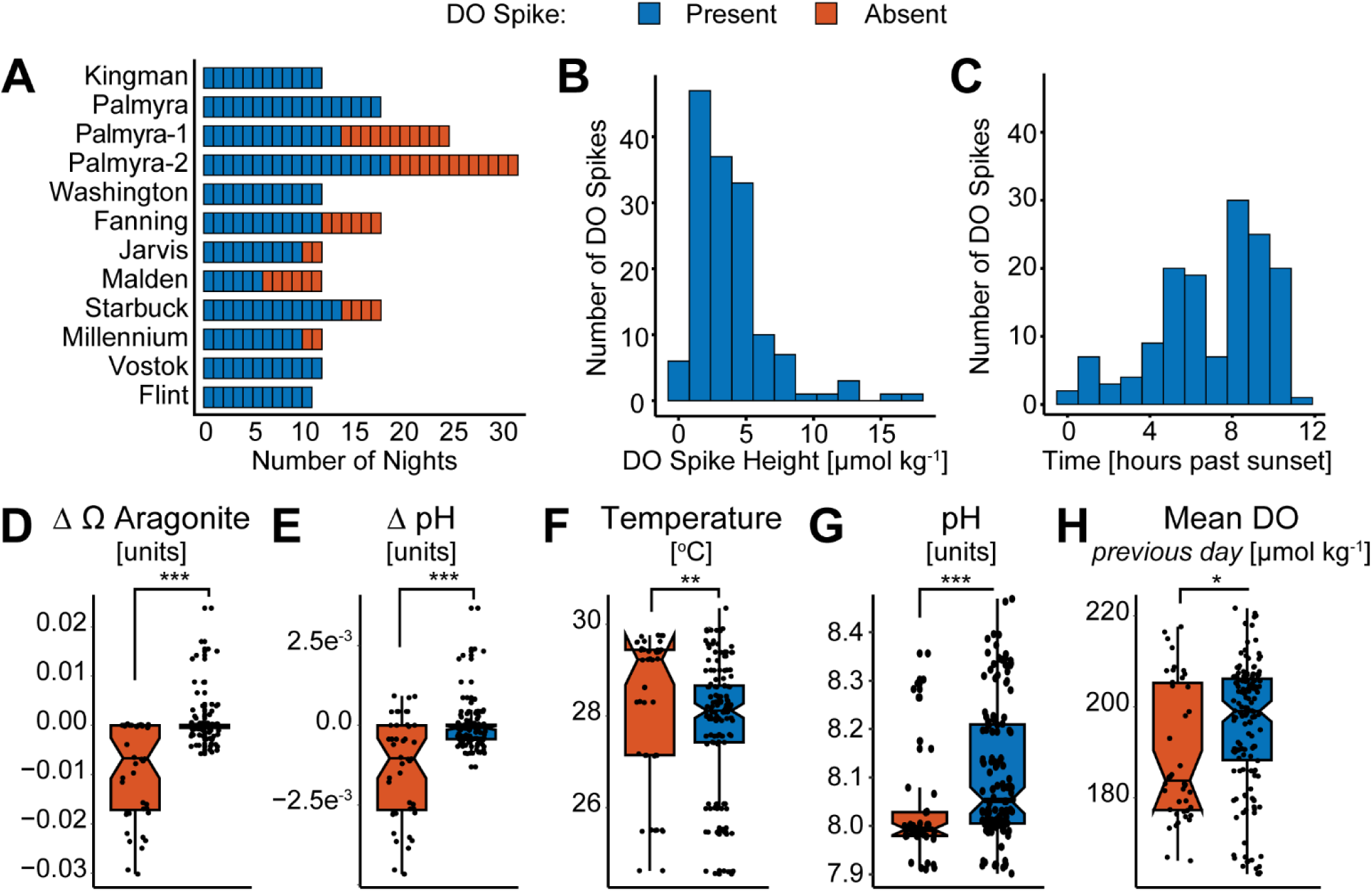
The saturation state of aragonite (Ω aragonite) is the most accurate predictor of nighttime dissolved oxygen spikes via random forests analysis. (A) Frequency of oxygen spike occurrence by night for each data set. ‘Palmyra-1’ and ‘Palmyra-2’are calcifier and non-calcifier dominated sites, respectively from the Palmyra BEAMS data, island names are from the Line Islands cBIT deployments. The y-axis is ordered by decreasing latitude from top to bottom. (B-C) Histograms of binned oxygen spike heights and times. (D-H) Top five predictors for all data sets in the Line Islands (cBIT deployments and BEAMS) using site level and island level variables, as determined by mean decrease in accuracy and p-value < 0.05. Points represent the value at the time of a DO spike (D-G), or the mean value for the previous daylight period (H). Asterisks indicate significantly different means as determined by a Wilcox test. Thresholds: *** = p < 0.001; ** = p < 0.01; * = p < 0.05; ns = not significant. Out-of-box classification error rate = 4.6%.

### Statistical Analysis of Potential Mechanisms

The top predictors of DO spike occurrence are a mix of biologically related variables (Δ Ω aragonite, Δ pH, pH, and the mean DO from the previous day) and one physical parameter (temperature) (Figure 2D-2H). These variables were the best predictors when assessing the data at both the island-plus-site and site-only levels. The variation in each of these predictors between DO spike presence and absence helps describe what events are taking place when a DO spike occurs.

Biological variables show that Δ Ω aragonite and Δ pH are both near zero to slightly positive when a spike occurs, while pH is higher (Figure 2D, 2E and 2G). This indicates Ω aragonite and pH are approaching a local maximum, which can be seen in the described behavior for DO spike presence and absence constructed using the original values plus first and second derivatives (Figure S6H and S6L). For the combined datasets pH values are higher when DO spikes occur but not Ω aragonite (Figure 2G and S6I). However, when comparing each dataset individually, it becomes apparent that the Ω aragonite is also higher when DO spikes occur (Figure S7A and S7D), but this pattern is lost when combining the Line Islands cBITs and Palmyra BEAMS data. Each dataset individually shows that Ω aragonite is higher when DO spikes occur, but the Line Islands cBITs data has overall lower Ω aragonite values than the BEAMS data. The other biologically relevant parameter from the random forests analysis describes DO concentration during daylight hours, where the mean DO from the previous day is higher when a DO spike occurs (Figure 2H).

Two additional biological variables exclusive to the Palmyra BEAMS dataset (and thus excluded from the random forests analysis) are derived from DO and pH data: net community production (NCP) and net community calcification (NCC). Neither of these have different original values between when a DO spike occurs and when it does not (Figure S6M and S6Q), but Δ NCC and ΔΔ NCP are both significantly different (Figure S6N and S6S). These describe at pattern where NCC is flat and NCP is at a local minimum when DO spikes occur, while NCC is steadily dropping and NCP is flat when DO spikes do not occur (Figure S6P and S6T). Temperature is lower when DO spikes occur (Figure 2F) and is stable, while it is higher but steadily dropping when DO spikes do not occur (Figure S7H). One island level physical variable, hydrodynamic pressure at 300 m depth was selected as a significant predictor during random forests analysis, but no significant differences were seen between DO spike presence and absence (Figure S7A – S7D).

Several mixed-effect linear models using the variables identified in the random forests analysis and combined to form a structural equation model (SEM) (Table S5) confirm that the strongest predictors of DO spike occurrence are pH, Ω aragonite, and temperature (Figure 3A). The overall strongest predictor of spike occurrence is the actual DO concentration at the time of a spike (coefficient: +33.64, p-value: 0.0012, Table S6), indicating that overall nighttime DO is not so variable as to prevent spikes from consistently producing DO concentrations higher than the DO concentrations when spikes do not occur. The second strongest overall predictor of DO spikes is ΔΔ pH (coefficient of −33.28, p-value: 0.0032), confirming that pH does indeed reach a local maximum and not a minimum alongside DO. The first derivative of DO directly correlates with DO spikes, so this relationship is heavily penalized by the correlation structure in the model and does not show up in the SEM plot. However, predictors of Δ DO can generally be assumed as predictors of DO spike occurrence since the model for each of these variables is effectively predicting the same event using either regression or binary classification, respectively (Table S5). Temperature is a strong predictor of Δ DO (coefficient: −22.67, p-value: 0.001), which matches the finding that temperature is lower when DO spikes occur. The nature of the relationships between pH, Ω aragonite and DO spikes are also validated by the SEM analysis, where Δ pH and Δ Ω aragonite are positive predictors of DO spikes and Δ DO, indicating that they each reach local maxima together.

**Figure 3.**
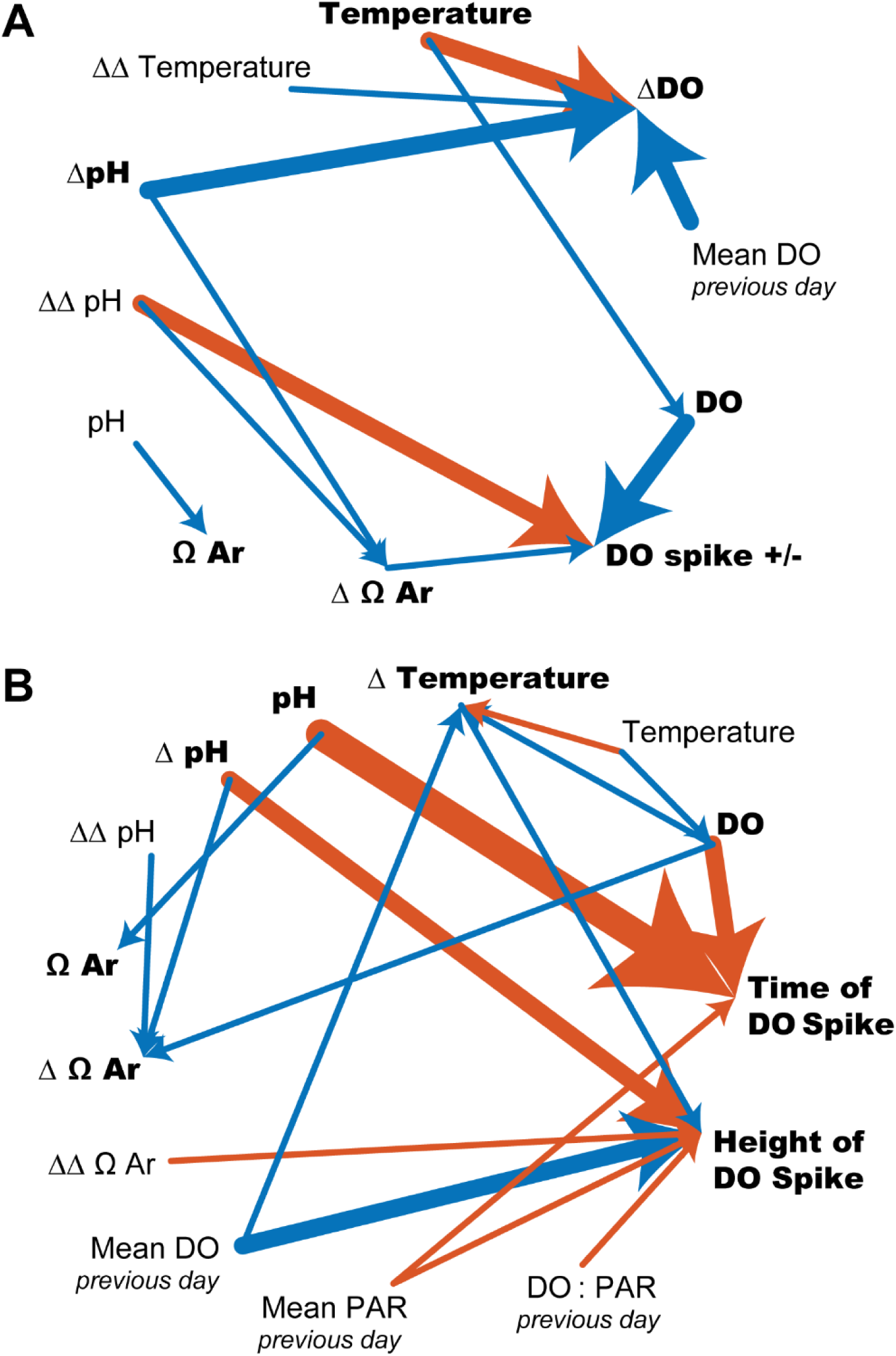
Omega aragonite, pH and temperature all interact to predict DO spikes, excluding most other modeled variables. The response variables for each model are indicated in bold. Relationships with a p-value < 0.05 as determined by Kenward-Rogers approximation and an SEM coefficient >|1| are shown. Arrows point from predictor to response. Thicker lines indicate stronger relationships as determined by the absolute value of the SEM coefficient. Orange lines indicate a negative effect, blue indicates positive. (A) Classification of DO spike presence/absence. (B) Prediction via regression of DO spike height and time

Another set of nested linear models using only data where DO spikes are present (Table S7) shows that in addition to pH and Ω aragonite, the levels of DO and PAR present the day before a DO spike occurs can predict the height of the spike (Figure 3B). The strongest predictor of either height or time is pH, which is a very strong predictor of DO spike time (coefficient: - 124.84, p-value: 0.006, Table S8). The strongest predictor of height is Δ pH (coefficient: −15.39, p-value: 0.0001). Mean DO concentration the day before a DO spike occurs is the second strongest predictor of spike height (coefficient: +13.33, p-value: 0.023). This also confirms the random forests prediction of higher mean DO the day before a DO spike occurs. Overall these data reinforce the significance of the positive relationship between pH and nighttime DO spikes.

Robust regressions (regressions that iteratively weight the median data values higher than the upper and lower extrema) between selected variables of interest from analyses up to this point elucidate a more nuanced relationship between nighttime DO spikes and pH related parameters (Figure 4). The sums of nightly NCC and NCP (ΣNCC and ΣNCP) for both sites in the Palmyra BEAMS dataset show different relationships to DO spike height between the two sites (Figure 4A & 4B). ΣNCC becomes more positive, indicating more net accretion of calcium carbonate/calcification as DO spike height increases at the calcifier-dominated site (purple solid line - Figure 4A). ΣNCP on the other hand, becomes more negative indicating increased respiration (purple solid line – Figure 4B). On the non-calcifier dominated site relationships between ΣNCC or ΣNCP were not observed (green dotted line – Figure 4A and 4B). DO and Ω aragonite at the time of a DO spike for these two sites also do not have a significant linear relationship to spike height, showing that the instantaneous values for these two parameters are not drivers of height in this dataset (Figure 4C & 4D). However, significantly positive relationships between DO and Ω aragonite to spike height are present in the combined Line Islands cBITs and BEAMS datasets (Figure S11). Additionally, positive coupling exists between temperature and spike height, and temperature and pH, while there is a negative association between pH and the time an DO spike occurs (Figure S12A – S12C). This demonstrates that increasing temperature leads to larger DO spikes, while spikes happen later when pH is lower. The mean DO concentration the day before a DO spike was not found to significantly increase or decrease in relation to spike height (Figure S12D), casting doubt on the predictive strength of this relationship.

**Figure 4.**
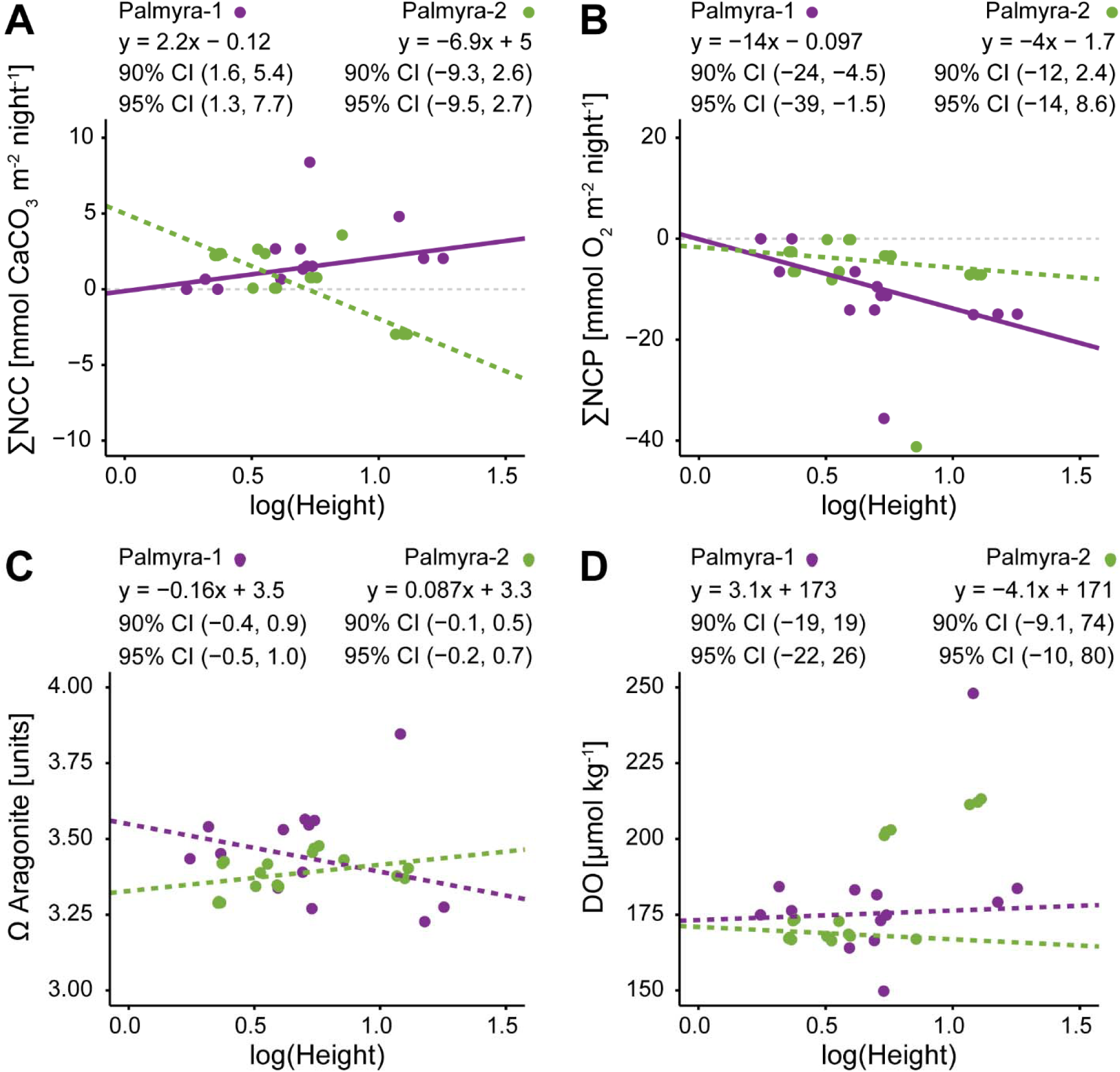
Net community calcification increases as oxygen spike height increases at a calcifier-dominated site (Palmyra-1), while the opposite is true for net community production. Robust regression analysis of log transformed oxygen spike heights and (A) nightly sums of NCC, (B) nightly sums of NCP, (C) saturation state of aragonite, and (D) oxygen concentration for the Palmyra BEAMS dataset. Dotted lines indicate non-significant regressions due to the inclusion of zero in both confidence intervals.

One physical variable, hydrodynamic pressure at 300 m depth had a significant correlation with DO spike height, while site level physical variables (observed in the Palmyra BEAMS data) show no linear relationship to height (Figure S13). However, there are significant differences between when a DO spike occurs and when it does not for the site level variables (Figure S8I – S8T). The combination of these findings indicates that locally pressure is higher when DO spikes occur and increases offshore as spike height increases, which describes a relatively calmer hydrodynamic environment. The observation of a higher percentage of nighttime DO spikes in the semi-enclosed cBITs also supports a positive relationship between these spikes and comparatively slower water movement. However, local current speed and directional shifts from the BEAMS dataset are associated with times when DO spikes occur, which in total describes a situation of calmer hydrodynamic conditions becoming less so.

Up to this point, all analyses presented here have been carried out on data representing only the exact time of a nighttime DO spike, excluding the extensive information contained in the rest of the time series data. Four separate generalized additive models (GAMs) built using site level physical variables, island level physical variables, biologically mediated variables, or a combination of all variables for the entire Palmyra BEAMS dataset show that NCC, NCP and Ω aragonite plus local current data are best at predicting if a DO spike will occur (Figure S14). The GAMs were most accurate when only nighttime data and only data collected closest to the benthos (for vertical gradient variables, see Methods) were used. However, this presented a problem when testing the models since subsetting training and testing data using very few DO spike observations was difficult. The GAMs fitted back to the original data do show that the combined GAM explains 81% of the variance in the data, which is an objectively good fit (Figure S14A). This validates the findings from all other analyses that drivers of nighttime DO spikes are predominantly related to NCC and NCP, with additional influence from water currents.

### In Vitro Isolation of Nighttime DO Spikes

A multitude of different incubation experiments were performed under controlled light and temperature conditions to determine if nighttime DO spikes could be isolated from open ocean influences (Figure 5, Tables S9 – S11). In total, 68 separate incubations were carried out, with 125 observations among them due to some incubations lasting several diurnal light cycles (Table S9 and S11). Of those observations, nighttime DO spikes occurred 23% of the time (29 out of 125). Crustose coralline algae (CCA, wild collected, Figure5H and 5I) incubations produced DO spikes at a rate of 37% (16 out of 27), and coral (aquarium cultured *Montipora capricornis*, Figure 5G & 5J) incubations at a rate of 32% (12 out of 25). DO spikes did not occur with incubations of other benthic organisms/samples after correcting for diffusion across the air-water interface in all non-sealed incubations (Figure 6A, Table S10). DO spikes occurred most frequently between 12 hours and 2 days of incubation time and were generally between 1.0 and 3.0 µmol kg^-1^ in height, with a few exceeding 30.0 µmol kg^-1^ (Figure 6B and 6C).

**Figure 5.**
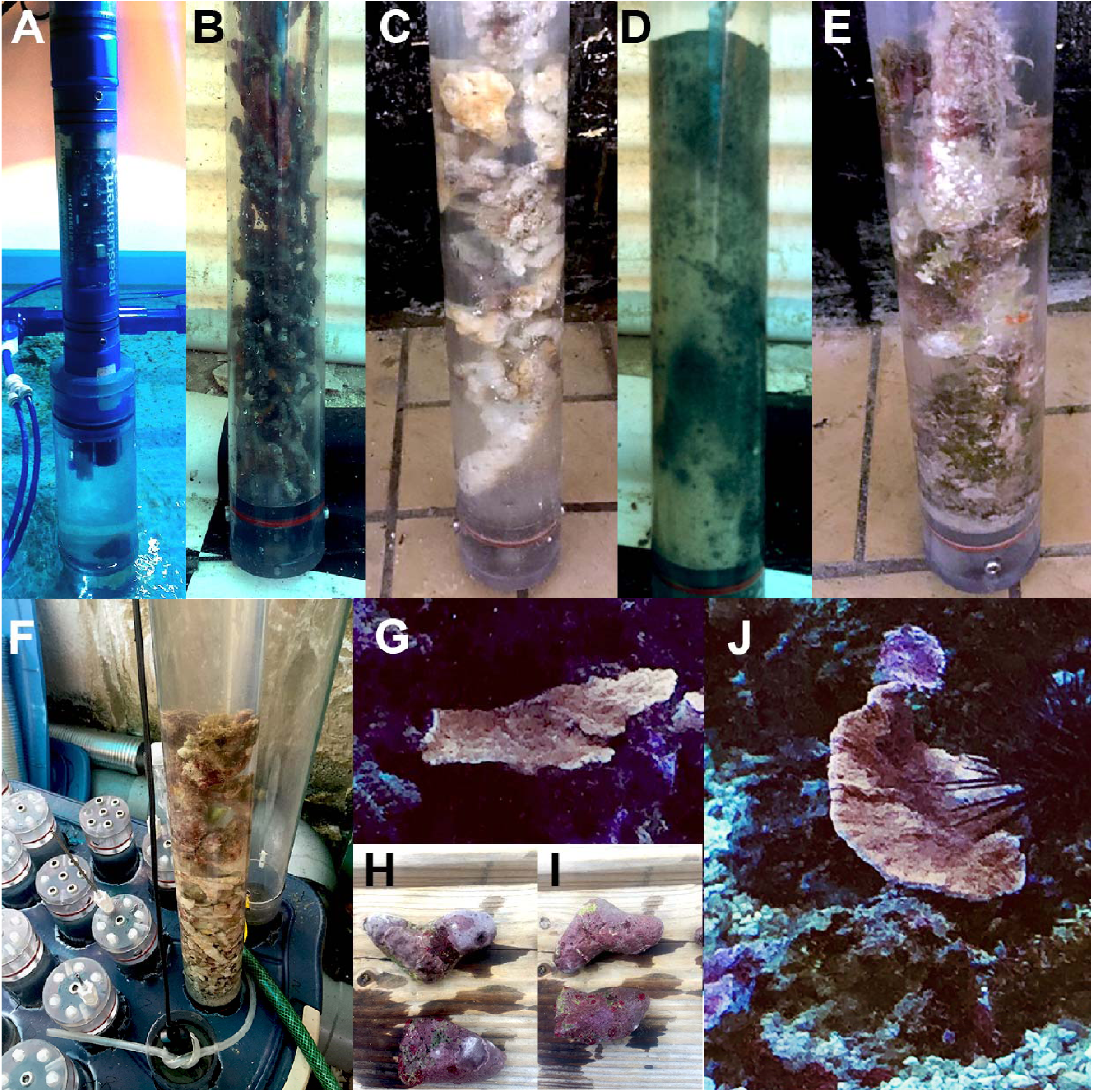
Controlled incubations were performed using wild collected and aquacultured reef organisms. (A) Position of MANTA sonde in fully sealed incubations. The incubation chamber pictured is smaller than those used in this study. (B – E) Wild collected turf, bare rubble, sediment and turf samples. (F) Layout of incubation chambers in water bath showing lids with access ports and a HACH probe in use (chamber without lid at bottom). (G & J) Aquacultured *Montipora capricornis* specimens before removal for incubation, appox. 10 cm long, 7 cm wide and 0.5 cm thick each. (H & I) Wild collected CCA specimens from intertidal zone, appox. 5 cm long, 3 cm wide and 2 cm thick each.

**Figure 6.**
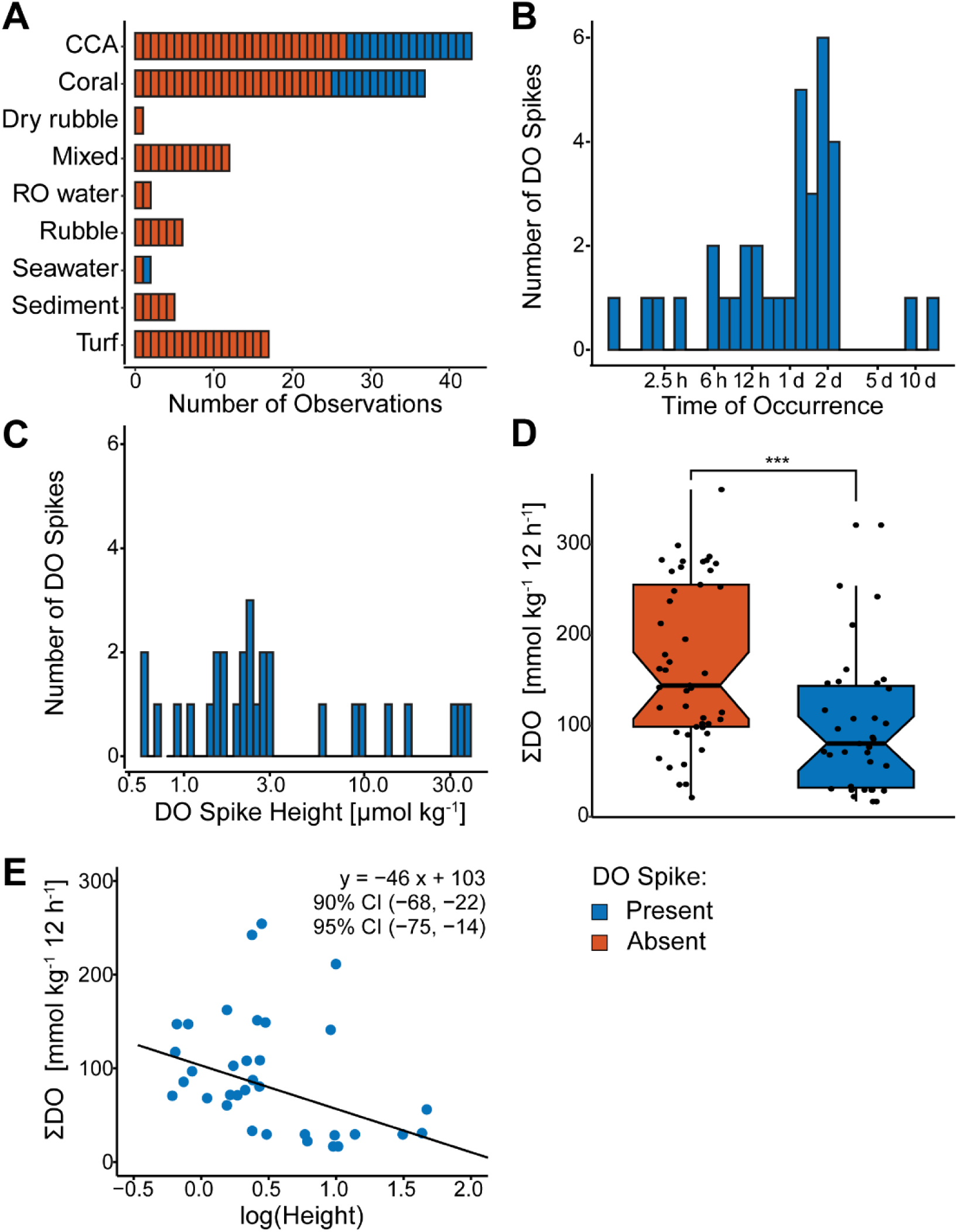
Analyses of incubation data show net community production (as total oxygen production) decreases as oxygen spike height increases in controlled incubations. (A) Frequency of oxygen spike occurrence for each organism incubated. (B-C) Histograms of binned oxygen spike heights and times. (D) Asterisks indicate significantly different means as determined by a Wilcox test. Thresholds: *** = p < 0.001; ** = p < 0.01; * = p < 0.05; ns = not significant. (E) The linear formula obtained by robust regression is listed on each plot, as well as the 90% and 95% confidence intervals obtained via bootstrap of robust regression.

The sum of oxygen present per 12-hour period can reasonably be assumed to represent ΣNCP per night in these incubations as any non-biological oxygen inputs have either been eliminated (by fully sealing the incubation chamber in an air-tight manner) or accounted for (by subtracting the amount of DO due to passive diffusion for open incubations). The ΣNCP is significantly lower when DO spikes occur in incubations and decreases as spike height increases (Figure 6D and 6E). These findings show that DO spikes can occur when fully isolated from non-biological variables and spike height increases with increasing respiration, as seen with the Palmyra BEAMS calcifier-dominated site. The observation of nighttime DO spikes in incubations of CCA and coral, but no other benthic organisms also supports the relationships between NCC, NCP and DO spike height observed at the calcifier-dominated site.

## Discussion

Here we provide unequivocal evidence that nighttime DO spikes occur regularly on the coral reef benthos around the world. We used a combination of peer reviewed, published data from 1970-present, our own field-based data collected from two ocean basins and data collected in controlled laboratory settings to corroborate these findings. In addition to documenting the existence of nighttime oxygen spikes in the environment, we sought to identify potential causal mechanisms. After extensive analyses of a variety of physical, environmental and oceanographic predictor variables, we were able to identify pH and Ω aragonite as the strongest predictors of DO spike occurrence and describe a positive correlation between calcification (ΣNCC per night) and DO spike height alongside a negative correlation between productivity (ΣNCP per night) and spike height. Further, detailed laboratory studies under controlled conditions allowed our team to isolate the pattern in a closed system, suggesting that DO spikes are likely the results of a biological, rather than a physical, process.

While it is obvious that physical forces influence DO concentrations during the night on reefs, the data presented here do not suggest that abiotic mechanisms are exclusively responsible for the existence of nighttime DO spikes. Abiotically, DO does not directly influence pH because it does not react with water to form H^+^. While DO is correlated with pH in marine systems due to biological influences (e.g., photosynthesis and respiration), physical processes transport water that shows the same correlation for the same reasons. This may partially explain pH and Ω aragonite as drivers for DO spikes, however the SEM approach used in this study is specifically designed to describe directional, causal relationships (Shipley, 2009, 2013). No significant relationship where DO causes pH changes was found, whereas pH was strongly predicted to cause all aspects of DO spikes. Δ pH slightly decreases from zero as DO spike height increases (Figure S11B) in a relationship shown to be a strong predictor of height (Figure 3B). This indicates that as DO spikes get larger pH reaches a local maximum just before DO does, which does not support a situation where DO and pH always change simultaneously. DO spikes causing pH spikes also does not explain nightly ΣNCC increasing as the height of DO spikes increases at a site dominated by calcifying organisms, since Ω aragonite (and thus pH) were shown to be decoupled from increasing spike height at this site (Figure 4). Takeshita et al. (2016) also found that daily ΣNCC was decoupled from daily mean Ω aragonite for both Palmyra BEAMS sites. Therefore, it is likely a more complex relationship is present during nighttime DO spikes then simply an external input of oxygen increasing pH and Ω aragonite and that subsequently increases NCC.

In order for oxygen to increase there must be either autochthonous oxygen (e.g. biological release *in situ*) or allochthonous oxygen inputs (e.g. transport by physical processes such as tides, currents or wind-driven waves), both of which have been addressed in this study. Our observation of increased DO during fully sealed incubation chambers provides evidence that these benthic organisms are capable of producing DO during the night. Physical contributions in these experiments from convection due to a temperature gradient (coupled to an oxygen gradient) in the vertical incubation chambers can be ruled out by the observations of some of the largest DO spikes in horizontally oriented, fully sealed chambers (Table S10). Further evidence in support of this can be seen in two of the highest DO spikes from incubations, each above 30 µmol kg^-1^ (Figure S16). These two incubations represent two different organisms (CCA and coral) in two different incubation setups (an open vertical chamber and a fully sealed horizontal chamber) (Figure S16A and S16B, respectively). The chamber incubated horizontally with the sensor positioned in the side and at the same level as the coral sample (also closer to, since the coral was positioned in the middle of the chamber) shows more evidence of DO spike correlation with temperature fluctuations than the vertical chamber. This is likely caused by on and off cycles of the climate control system in the room where the incubation was carried out as well as increasing metabolic activity with temperature, rather than convective or diffusive gradient shifts.

DO spike height increases as ΣNCP decreases at a reef site dominated by calcifiers and in incubations, lending support to these spikes being positively correlated with increased respiration. DO spikes also generally occur either during or just after a period of calm hydrodynamic conditions yet changes in those same conditions are not associated with DO spike height. Temperature does increase with spike height although it is generally lower overall than when no spikes are observed. While at least one *in situ* study suggests that decreasing water flow leads to overall decreased respiration and calcification at night (Shaw et al., 2014), this is attributed to less available oxygen and DIC leading to less consumption of both. In the case of nighttime DO spikes, conditions appear to converge on a combination of increased consumption but less water movement. These observations suggest that DO spikes result from a punctuated uptick in metabolic activity under conditions that are not conducive to supporting such activity.

### Potential Biological Explanations

Based on these findings, we hypothesize that some of the nighttime DO spikes are a biological response by benthic calcifying organisms to increasing respiration combined with reduced exposure to more oxygenated water. This results in intracellular hypoxia and significant H_2_O_2_ release, which is subsequently degraded into DO and H_2_O via ROS detoxification activities of pelagic and benthic organisms. Hypoxia response in eukaryotes is highly conserved and involves H_2_O_2_ production from superoxide via the electron transport chain (ETC) (Guzy & Schumacker, 2006). Oxygen sensing molecules in mitochondria switch the ETC into superoxide production mode when intracellular oxygen reaches a critical threshold but is not exhausted (Murphy, 2009; Cadenas, 2018). In this manner some oxygen is used to create the powerful signaling molecule H_2_O_2_, which then turns on hypoxia inducible factors (HIFs) that activate a metabolic shift leading to reduced oxygen consumption (Semenza, 2007; Smith, Waypa & Schumacker, 2017). Coral HIFs have recently been described and coral have been shown to survive low oxygen levels around 1 mg l^-1^ (equivalent to 31.3 µmol kg^-1^) for up to 72 hours (Vaquer-Sunyer & Duarte, 2008; Zoccola et al., 2017), implying some degree of conserved, long-term hypoxia response. As discussed in the introduction, coral have also previously been shown to rapidly release H_2_O_2_ in concentrations that could result in its breakdown to form DO spikes of the size shown here. Crustose coralline algae also produce H_2_O_2_ in response to hypoxic stress which is often detoxified by bromoperoxidase, a potent generator of singlet oxygen (another type of ROS) that rapidly decays to ground-state molecular oxygen (Everett, Kanofsky & Butler, 1990; Triantaphylidès & Havaux, 2009; Wever, Krenn & Renirie, 2018). A large H_2_O_2_ release as respiration drives oxygen concentrations inside the cells of coral or CCA below the signaling threshold for hypoxia is the most parsimonious biological explanation for the nighttime DO spikes presented in this study at this time.

Increased calcification could also be a response to hypoxia, as suggested by Wooldridge (2013) with reference to a theoretical coral glyoxylate pathway (Kondrashov et al., 2006) that would help coral cells detoxify increasing concentrations of acetate, built up through prolonged fermentative metabolism (Müller et al., 2012). Wooldridge hypothesizes that acetate once converted to oxalate is pumped into the coral’s extra-cytoplasmic calcifying fluid where it complexes with calcium ions to form calcium oxalate crystals. These crystals in turn become nucleation sites for nighttime calcium carbonate precipitation. While experimentally unverified as a complete process, several other studies have remarked on patterns of dark calcification and respiration similar to those in our study and speculated that a link between increased calcification and environmental stress may be at work (Domart-Coulon et al., 2014; Jokiel, Jury & Rodgers, 2014; Rippe et al., 2018). Calcifying red and green algae (e.g., CCA) are also known to use the glyoxylate pathway in a comparable manner to precipitate their calcified structures (Pueschel & West, 2007; Yang et al., 2015).

### Relevance to Coral Health

Nighttime DO spikes, given our hypothesis is further verified, are likely part of a hyperbolic response to hypoxia wherein the DO spikes occur at an early to intermediate stage of hypoxic stress and then give way to more drastic reductions in metabolic activity, similar to the general response curves described by Nelson and Altieri (2019). Increased respiration in at least one species of coral an hour after exposure to hypoxia has been reported, followed by respiration decrease after longer exposure (Dodds et al., 2007). Many studies of the effects of hypoxia on corals have used environmental conditions at less than 50% air saturation, more in line with the conditions in our incubations where DO often dropped well below 50% saturation, but with much greater water flow and only the differences in respiration or calcification from beginning to end points are reported (Al-Horani, Tambutté & Allemand, 2007; Wijgerde et al., 2014; Osinga et al., 2017). We see DO spikes *in situ* at DO concentrations that are never below 70% saturation, meaning that these DO spikes are perhaps one of the first environmental signs of intracellular hypoxia. Hypoxic stress is quickly gaining attention as an ongoing threat to coral reef health due to rising temperatures, eutrophication and competition with fleshy macroalgae (Barott et al., 2009; Haas et al., 2013b,a; Sugden, 2017; Roach et al., 2017; Gajdzik & DeCarlo, 2017; Breitburg et al., 2018; Nelson & Altieri, 2019). H_2_O_2_ in excess has also been linked to coral bleaching (Downs et al., 2002). If regular observations of DO spikes are an indicator of repeated but moderate hypoxic stress, they would be a sort of bellwether for greater susceptibility to coral bleaching events, disease and even mass mortality. It would be important to include them in future reef health assessment systems for this reason.

### Limitations of Statistical Inference

The goodness of fit data for the SEM analyses indicate that time was not well modeled at an r-squared value of 0.34 even after accounting for island as a random variable (Figure S10B). This is the lowest r-squared for any model in either SEM analysis and implies that additional data/modeling approaches may be needed to improve the fit. However, the p-value of 0.409 for the height-time SEM (Table S7) leads to rejection of the null hypothesis that there are significant relationships in the data not represented in the model. Thus, of all the relationships represented in the SEMs, relationships with time are the most questionable. The remaining relationships are well fit and trustworthy, even with a p-value of 0.004 for the DO spike occurrence SEM (Table S5). This is due to three significant relationships not included in the model because they do not make sense from a causal standpoint (spike occurrence as a predictor for Δ DO, and two aspects of DO as predictors of temperature). Likewise, the GAM analysis was incomplete due to limited observations of DO spikes in the Palmyra BEAMS data that made testing the model difficult. Yet, the combined model fits the data well allowing for some degree of trustworthiness to exist. Additional time series data of the type in this dataset should be used in future modeling efforts to strengthen the causal inferences.

There are more total observations for CCA and coral samples in the incubation data than any other organism or sample type. With a DO spike occurrence rate of only 23% it is tempting to assume that there were not enough incubations of other types for spikes to be observed. However, the first set of incubations done in 2015 included all wild collected benthic organism/sample types in equivalent numbers of incubations (see Methods). Once the largest DO spikes were observed in CCA samples and DO spikes in other samples were eliminated after correcting for diffusion in the open incubation chambers, subsequent incubations focused on CCA and later coral.

## Conclusion

This study clearly identifies the existence and prevalence of nighttime DO spikes from coral reefs around the world. Further, these data provide evidence that nighttime DO spikes are at least partially biological in origin and that this process has a significant effect on coral reef productivity, a finding that demands more research. Future studies should focus on further analyses of both *in situ* and *in vitro* data, especially mechanistic studies that reveal the source of these anomalies. Rigorous models based on statistical learning methods can be developed from additional time series data and at specific points when spikes occur to further evaluate the mechanism driving DO increases at night. Incubation studies utilizing sensors for H_2_O_2_ as well as pH and discrete TA and DIC measurements will help further understanding of the calcification and hypoxia link, as well as provide more data for a biological model of what might be occurring. These findings have important implications for biological feedbacks, benthic boundary layer dynamics, hypoxia, reef metabolism and overall coral reef health and resilience. We hope that these results motivate future research to help resolve this widespread and ecologically important phenomenon.

## Supporting information

all supplemental

## Acknowledgements

The authors would like to thank Dr. Barbara Bailey, professor of statistics at San Diego State University (SDSU) and members of the Biomath Group at SDSU for their guidance regarding appropriate statistical models for this work. We also thank Mark Hatay of SDSU for his assistance constructing custom incubation chambers, as well as staff and researchers affiliated with the CARMABI research station, the crew of the Hanse Explorer, the Nature Conservancy and the Palmyra Atoll Research Consortium.

