## Supplementary material for "Evidence for a biological source of widespread, reproducible nighttime oxygen spikes in tropical reef ecosystems has implications for coral health": all supplemental

**Supplemental Methods**

**Curacao 2015**

The island of Curaçao (12°7’N 68°56’W) is located approximately 64 km northeast of the Venezuelan coast on the southernmost edge of the Caribbean tectonic plate. It is a semi-arid island surrounded by fringing reefs, with greater coral diversity and coral coverage than much of the Caribbean. Research on this island was conducted out of the CARMABI Research Station from April 14 to May 28, 2015. A mean temperature of 26.7 +/- 0.1^o^C was collected as described for the previous islands. A single MANTA multiprobe sonde was deployed in Curaçao at 10m of depth approximately 200 m offshore of a desalinization plant located at 12^o^6’N, 68^o^57’W. The MANTA was set up to autonomously log parameters per the deployment for Mo‘orea and the Line Islands, with the exception that no cBITs were used. No PAR data was taken due to the lack of an appropriate PAR sensor during this expedition.

**Mo’orea 2011**

Sites at the island of Mo‘orea (17°48’S 149°84’W) were monitored from September 1 through 22, 2011. cBITs were deployed for 36 hours at 5 m depth on the back reef habitat of Mo‘orea for the purpose of isolating the benthic water column from the surrounding seawater, after the methods described by (Haas et al., 2013). Briefly, each cBIT contained a MANTA multiprobe sonde (Eureka Water Probes, www.waterprobes.com) with sensors measuring and logging pH, redox potential (ORP), conductivity, dissolved oxygen (O_2_), and temperature on 5 minute intervals.

**Predictors of Oxygen Spike Occurrence for Whole Time Series**

A spline-based modeling approach was used to assess the Palmyra BEAMS time series data set for predictors of nighttime oxygen spikes. The BEAMS data set was used because of the length and variety of time series variables it contained, including site level water current data and fluxes of two important biological variables: NCC and NCP. Three separate models were constructed from either biological variables (O_2_, pH, NCC, NCP and Ω aragonite), site level physical variables (local current speed, current direction, water pressure and temperature), or island level physical variables (offshore water pressure at 300 and 500 m, wind speed and direction, and moon phase) using generalized additive models and a binomial distribution (R package mgcv, function gam). These separate models were used to determine if any of the three sets alone could accurately model the occurrence of O_2_ spikes. Variables for each model (including first and second derivatives) were added one by one and compared via ANOVA, while optimizing the best fit (adjusted r-squared), maximum likelihood, and AIC values.

A combination of P-splines (penalized basis splines, or splines constructed with n-order polynomials), cubic splines (splines constructed with 3rd-order polynomials), and cyclical cubic splines (cubic splines that repeat a pattern) were used to fit individual variables, while tensor products were used to create a 2-dimensional surface that fits a combination of two variables. A value called gamma is used to increase or decrease the degree of smoothing a spline has compared to the original data, where values over 1 increase the smoothing, values below 1 decrease smoothing. Gamma values between 0.3 and 1.0 were used improve the fit of certain splines. The number of points a spline smooths across is set by the value of k in the model. The default values for k were used unless the function gam.check indicated that k was too small for a certain spline. Data were split into night and day times and by distance from the benthos for all gradient measurements (pH, O_2_, temperature, and site level physical variables). Factors for site (either calcifying or non-calcifying) and hour of the day were also used. The best three models were combined into one final model to determine what, if any, contribution each predictor could make to a final prediction of nighttime oxygen spike presence/absence.

**Additional Incubation Information**

Incubations were carried out over a 12-hour period for all samples taken in Curacao during 2015, with individual incubations designated by number. A numbering scheme of ‘0.00’ was used, where the number in front of the decimal designates the arbitrary sample number of an individual sample (reset for each round of incubations), and the number after the decimal indicates the incubation round (numbered continuously). After each incubation, crustose coralline algae (CCA) samples were placed in either an open plastic bowl in a low-light aquarium with seawater flow (samples from >5 m), or a polycarbonate incubation chamber with all ports open anchored in the intertidal zone in front of CARMABI. The CCA samples were allowed to recover for a minimum of 24 hours to ensure sample survival for ongoing incubations.

During the 2016 experiments CCA samples were collected from various sites around the island of Curacao at depths ranging from 0.5 – 15 m and held in a shaded aquarium with flow until processing and incubation. Processing to generate CCA slurry was carried out using a steel chisel to gently chip away an approximately 1mm deep layer of the surface material of seven randomly selected CCA samples, followed by rinsing the chiseled surface and surface material with 0.22 μm filtered seawater into a 50 ml sterile conical tube labeled ‘Endolith’. The remains of each sample were then crushed with a hammer on a clean, dry surface and the innermost crushed rock placed in 30 ml of 0.22 µm filtered seawater in a 50 ml conical tube labeled ‘Endolith’. The CCA samples were split into two Endolith and five Epilith replicate incubations using 20 ml and 10 ml of each sample, respectively. This was done to balance the presumably lower biomass of the endolith sample against the higher biomass epilith. Three of the five epilith replicates were incubated in 1.5 l chambers with lids custom fit for use with a MANTA sensor (only three such incubation chambers were available) while all other incubations were carried out in 1 l chambers (measured by PreSens optodes). Each chamber was filled with 0.22 µm filtered seawater before inoculation with a respective sample, and then sealed with the sensor/lid combo, making sure to eliminate any headspace in the chamber. Incubations ran for 48 hours, allowing two full diurnal cycles to take place.

**Calculation of Diffusion for Open-Top Incubations**

Fick’s first law of diffusion is written as:

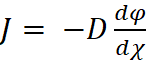

Where J is the diffusion flux, D is the diffusion coefficient, dφ is the change in concentration, and dχ is the change in height. For the open-top incubations, D was calculated from the salinity and temperature at each time point using Matlab function gas_diffusion and constants from Ferrell & Himmelblau (1967) and dφ was taken as the oxygen concentration at 100% air saturation minus the concentration measured. The change in height was assumed to be 1 cm for estimating the oxygen at the surface of the air-water interface, where the oxygen sensor was positioned. J in µmol O_2_ cm^-2^ s^-1^ for each time point was then multiplied by the surface area of the open top (33.1 cm^2^) and the time elapsed between each measurement (60 s), and then this value was subtracted from the observed oxygen measurement. This is likely an overestimate of the amount of oxygen present due to diffusion at each time point, since some of the surface area was occluded by the sensor. However, this further supports the magnitude of oxygen spikes observed after accounting for diffusion as not being driven by physical forces.

**Supplemental Figures**

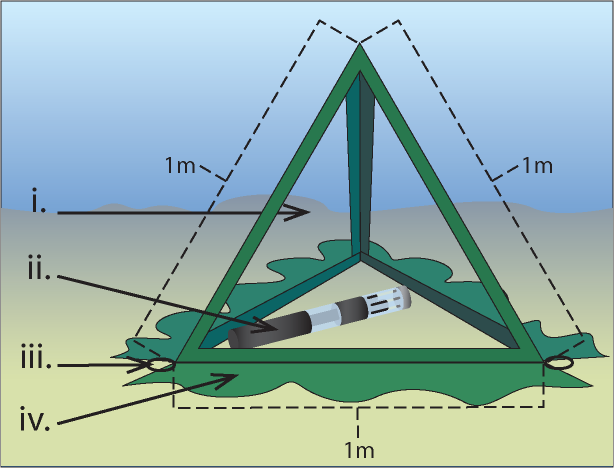

**Figure S1**. **Collapsible benthic isolation tent (cBIT) setup used for the Line Islands and Mo‘orea deployments.**

Described previously in Haas et al. (2013).

i. Transparent plastic window (polycarbonate)
ii. MANTA multisensor sonde
iii. Steel anchor rings
iv. PVC fabric skirt weighted to diminish flow into and out of cBIT.

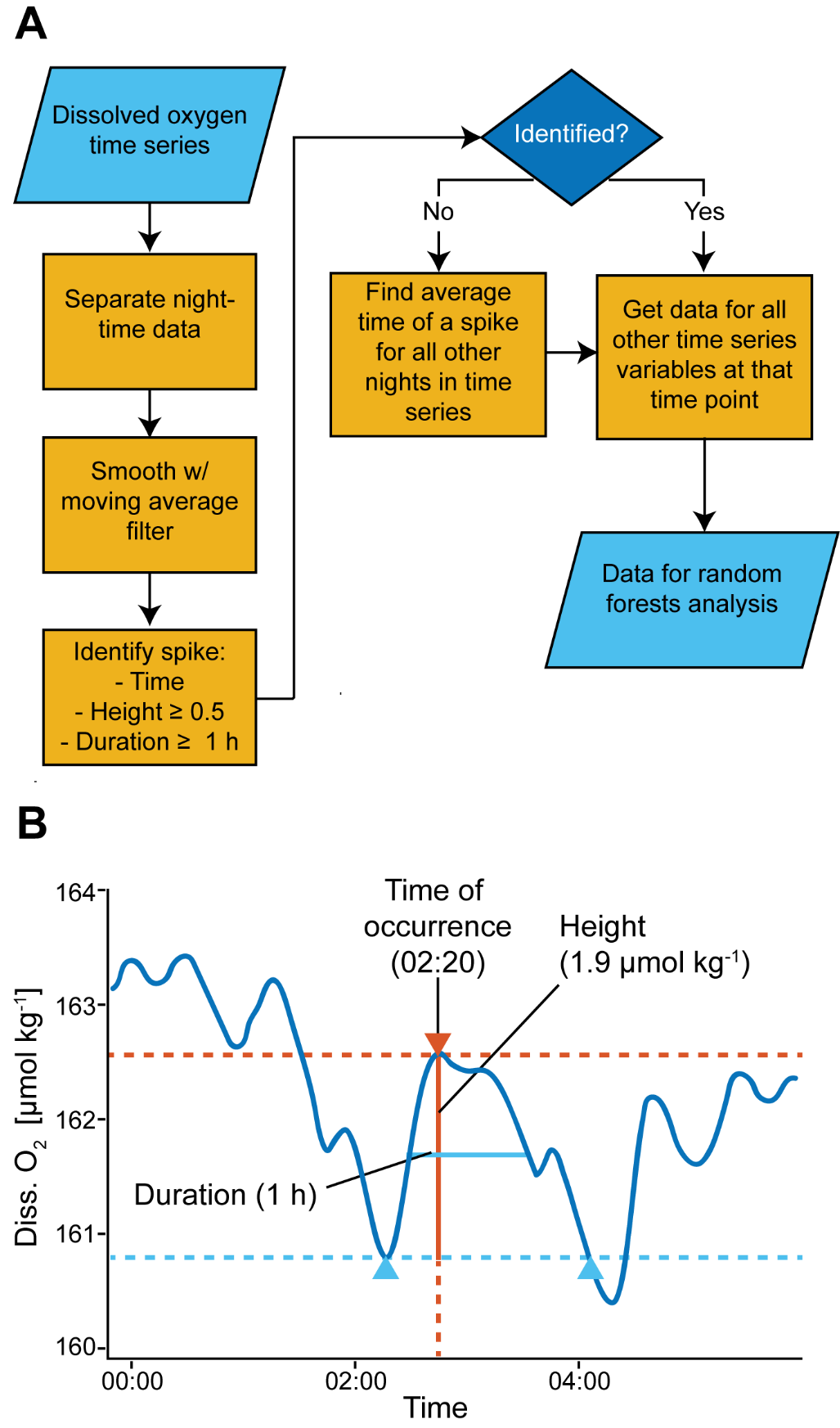

**Figure S2. Algorithm for identifying nighttime dissolved oxygen spikes.**

(A) Flowchart of algorithm steps applied to each time series dataset. (B) Illustration of the attributes used to identify a dissolved oxygen spike. Light blue dotted line indicates the baseline drawn between the two endpoints (light blue triangles). The width (duration – solid light blue line) is drawn at half the height (solid orange line). Orange triangle indicates the maximum value of the spike, which also determines the time of the spike.

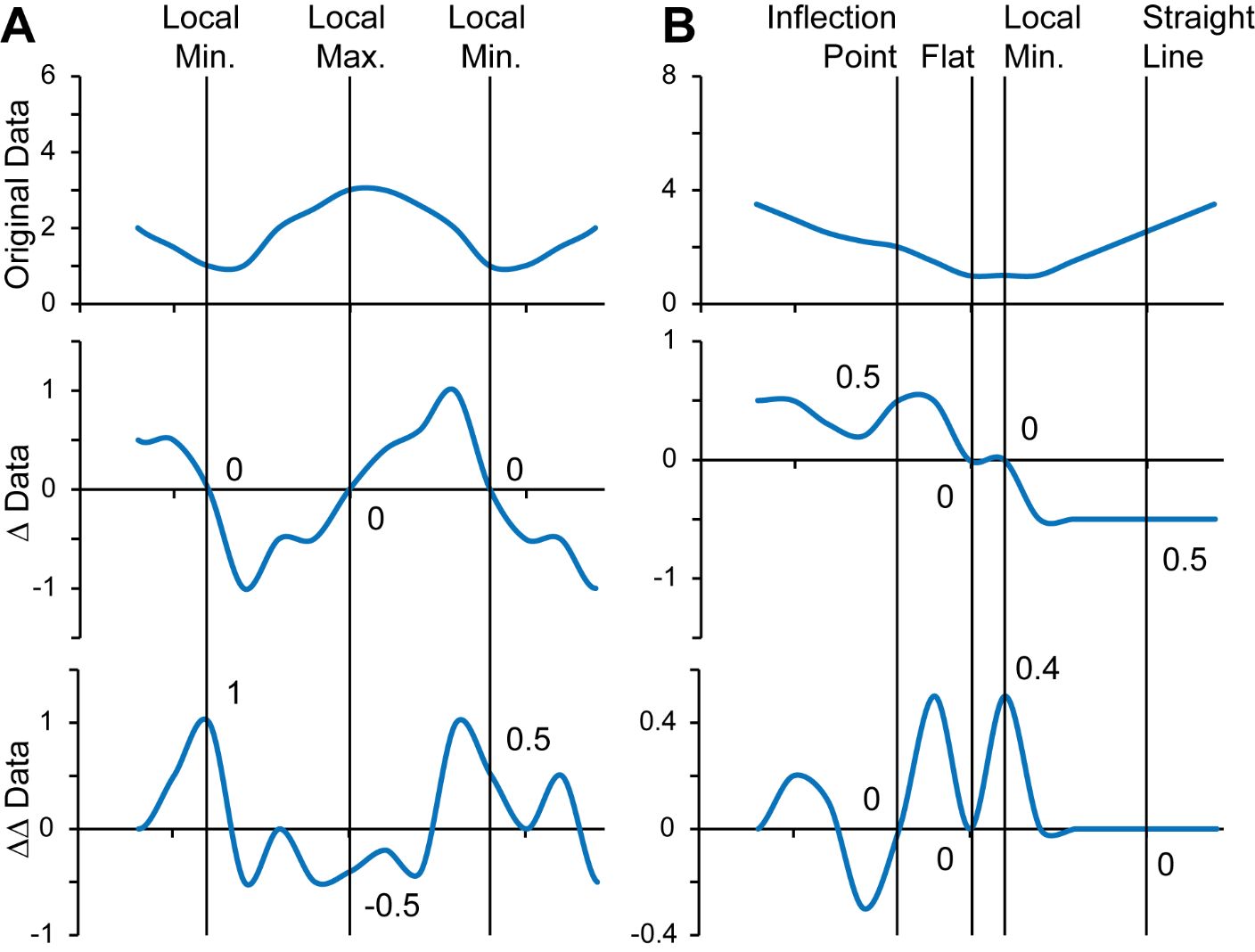

**Figure S3. The values of the first and second derivatives describe the behavior of a time series with respect to local minima, maxima and inflection points.**

(A) An example of the first and second derivatives of a time series similar to O_2_ measurements when an O2 spike occurs. (B) The same as A, but in the absence of an O_2_ spike.

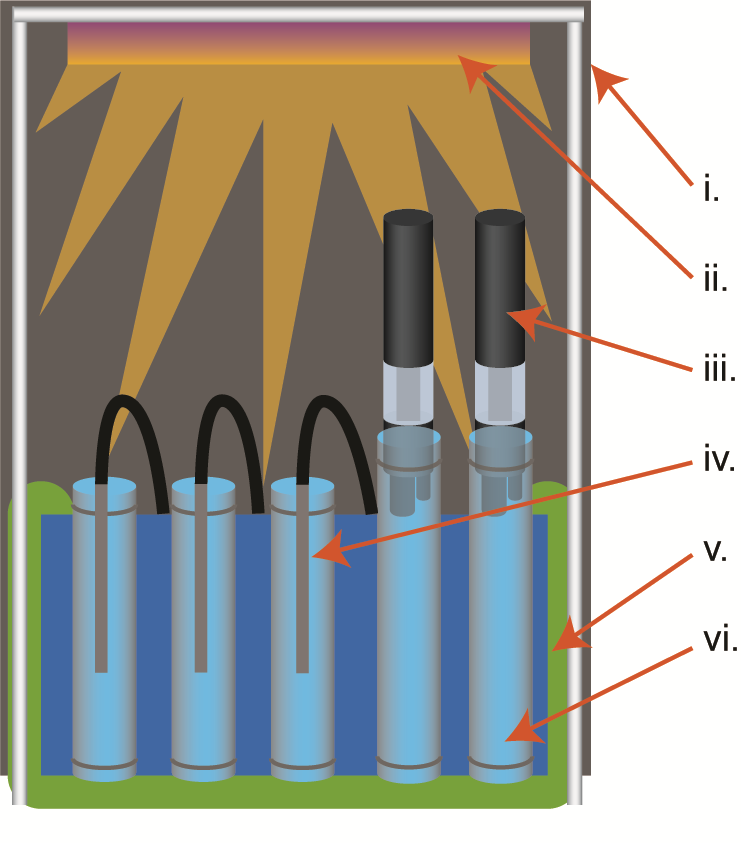

**Figure S4**. **Block diagram of laboratory incubation setup used in Curacao.**

i. Light-blocking fabric tent over PVC frame
ii. Aquarium lights controlled by a digital timer
iii. MANTA fitted into sealed 1.5 L inucbation chamber
iv. Fiber optic optode fitted into sealed 1 L incubation chamber
v. Temperature controlled water bath
vi. Location of intact samples

Note that the setup used between 2015 and 2016 was very similar, except during 2015 all incubation chambers were 2 L, incubations were only partially sealed, and no light was present.

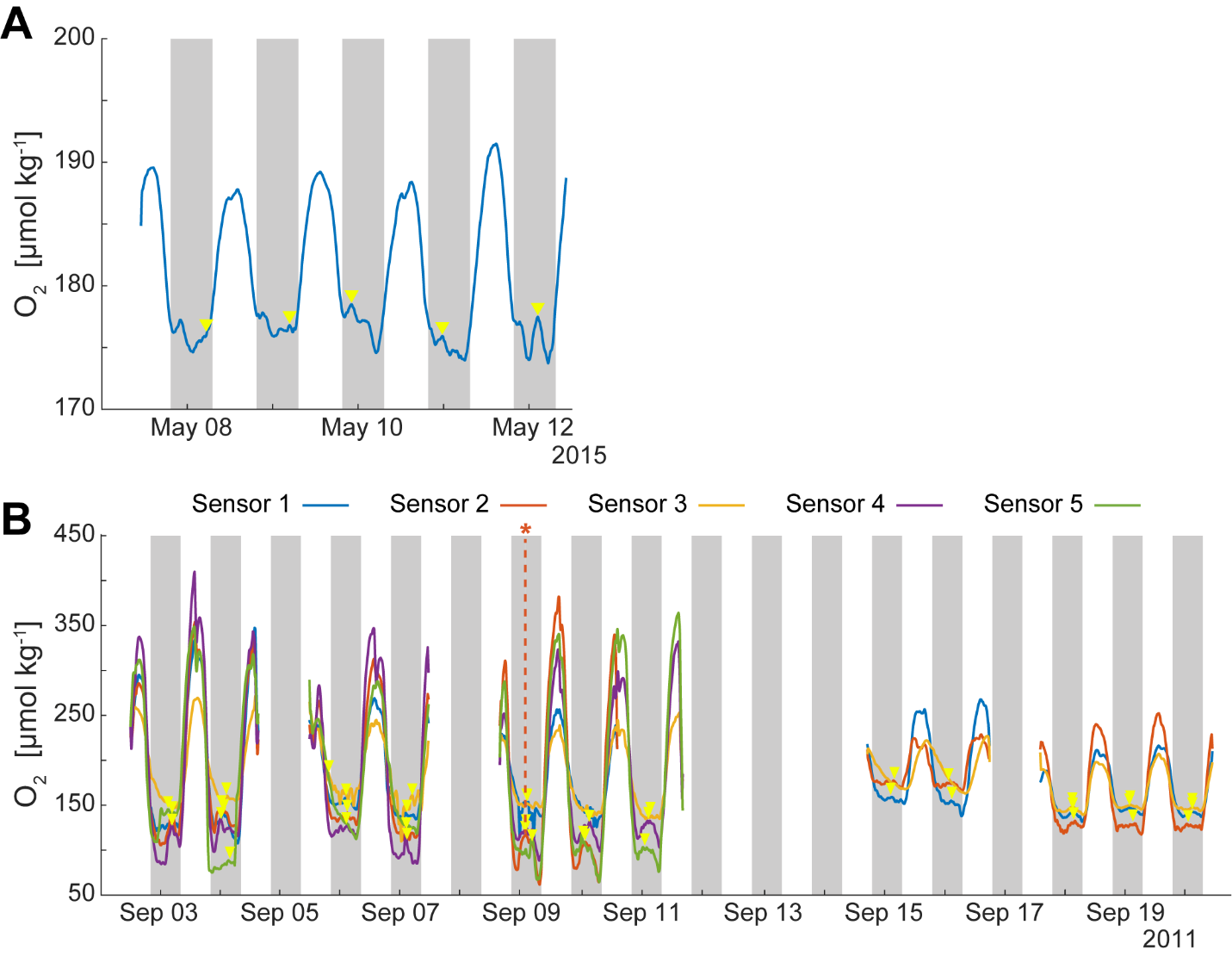

**Figure S5**. **Oxygen time series plots for Curacao and Mo’orea**

(A) Curacao – yellow triangles indicate the location of an oxygen spike. (B) Mo’orea - yellow triangles indicate the location of an oxygen spike, dotted orange line and asterisk indicate the tallest oxygen spike observed in any dataset.

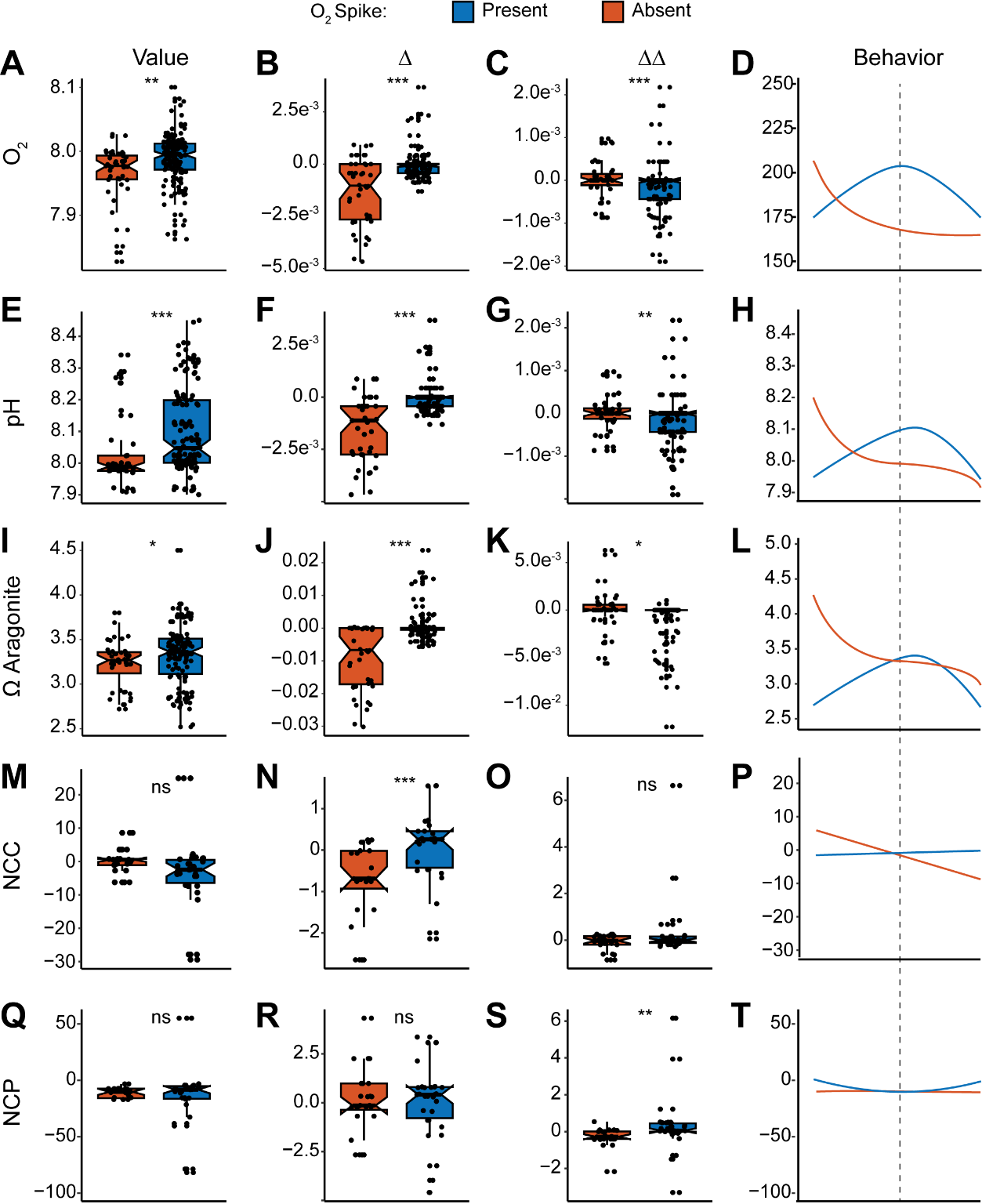

**Figure S6.** **Box plots of the first and second derivatives of significant biological variables at the time of an O_2_ spike, alongside the original values and the time series behavior the derivatives describe.**

Dotted line under Behavior column indicates the exact time of an oxygen spike. Asterisks indicate significantly different means as determined by a Wilcox test. Thresholds: *** = p < 0.001; ** = p < 0.01; * = p < 0.05; ns = not significant.

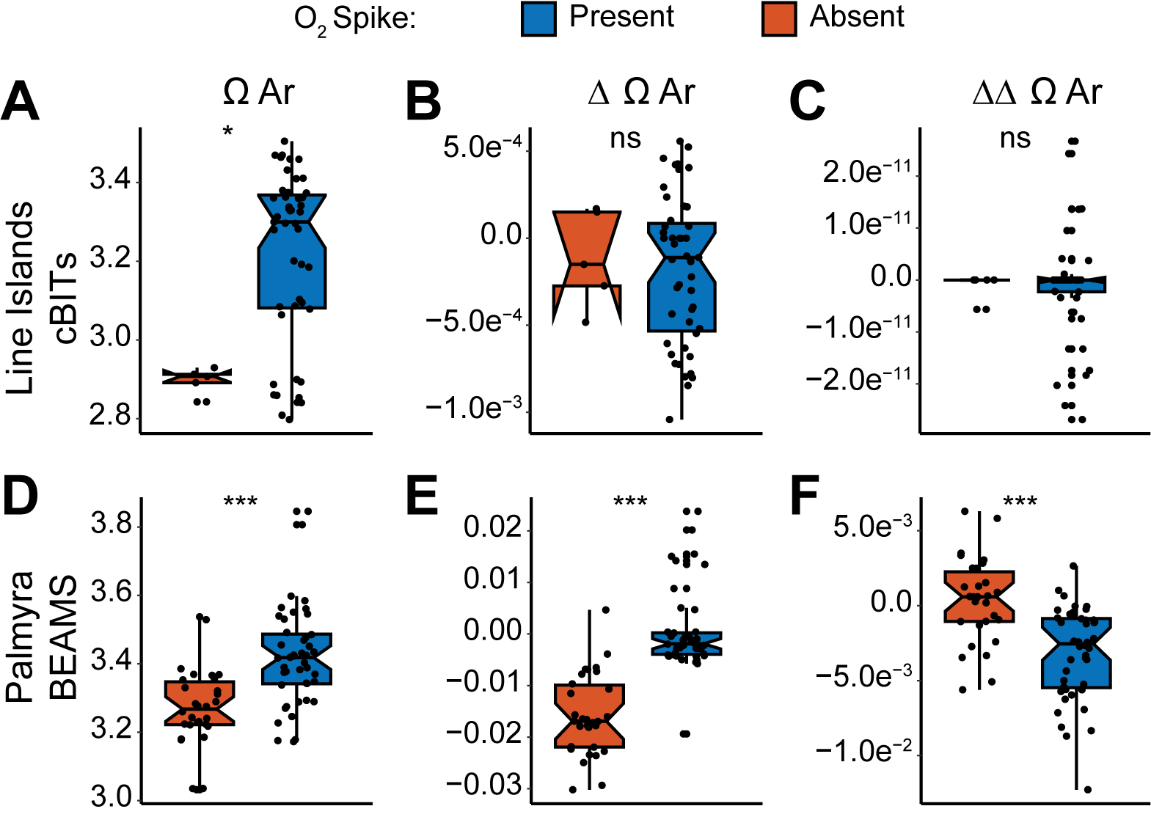

**Figure S7.** **Box plots of the first and second derivatives of significant biological variables at the time of an O_2_ spike, alongside the original values for omega aragonite between the Line Islands cBITs and Palmyra BEAMS datasets.**

Dotted line under Behavior column indicates the exact time of an oxygen spike. Asterisks indicate significantly different means as determined by a Wilcox test. Thresholds: *** = p < 0.001; ** = p < 0.01; * = p < 0.05; ns = not significant.

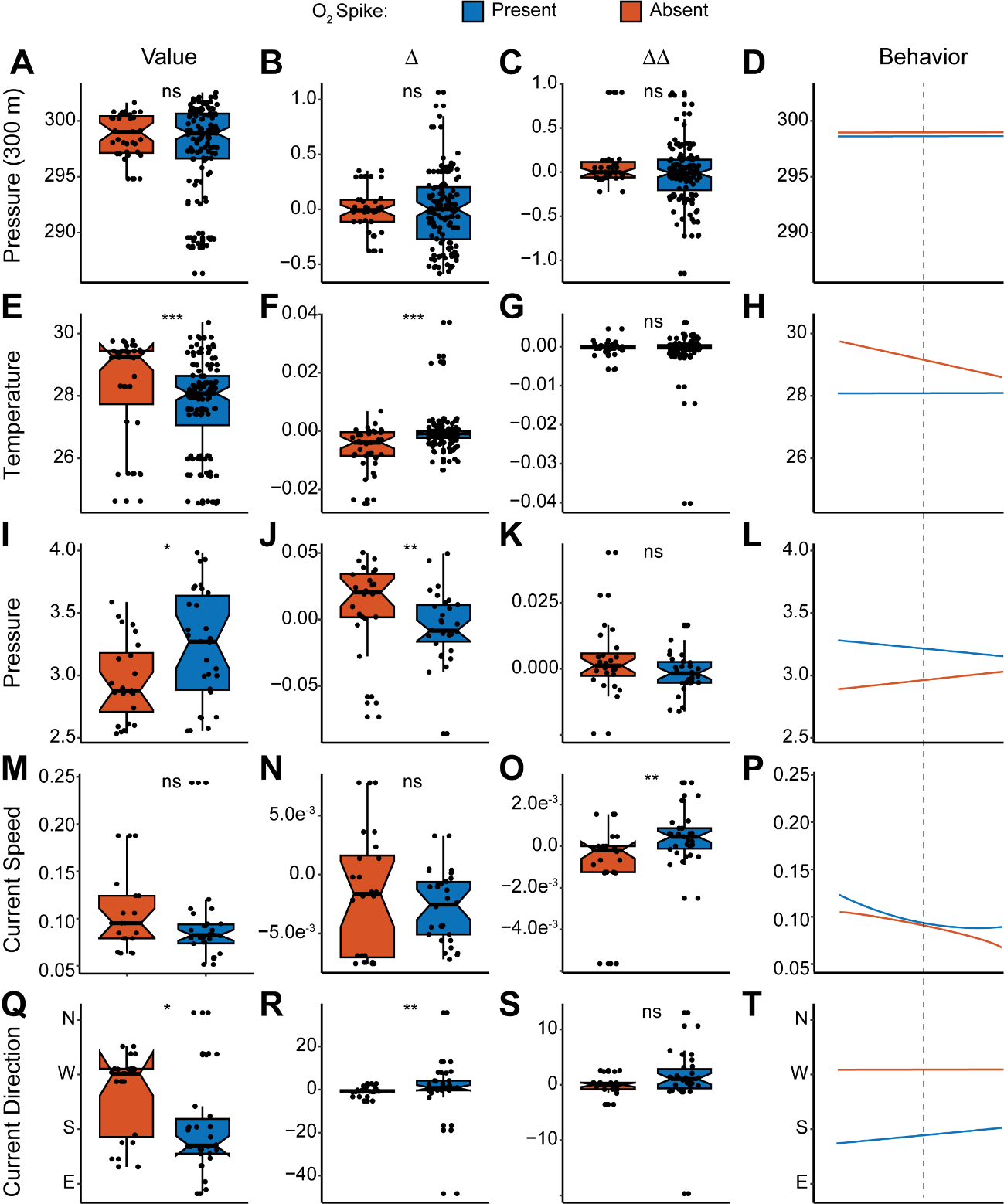

**Figure S8.** **Box plots of the first and second derivatives of significant physical variables at the time of an O_2_ spike, alongside the original values and the time series behavior the derivatives describe.**

Dotted line under Behavior column indicates the exact time of an oxygen spike. Asterisks indicate significantly different means as determined by a Wilcox test. Thresholds: *** = p < 0.001; ** = p < 0.01; * = p < 0.05; ns = not significant.

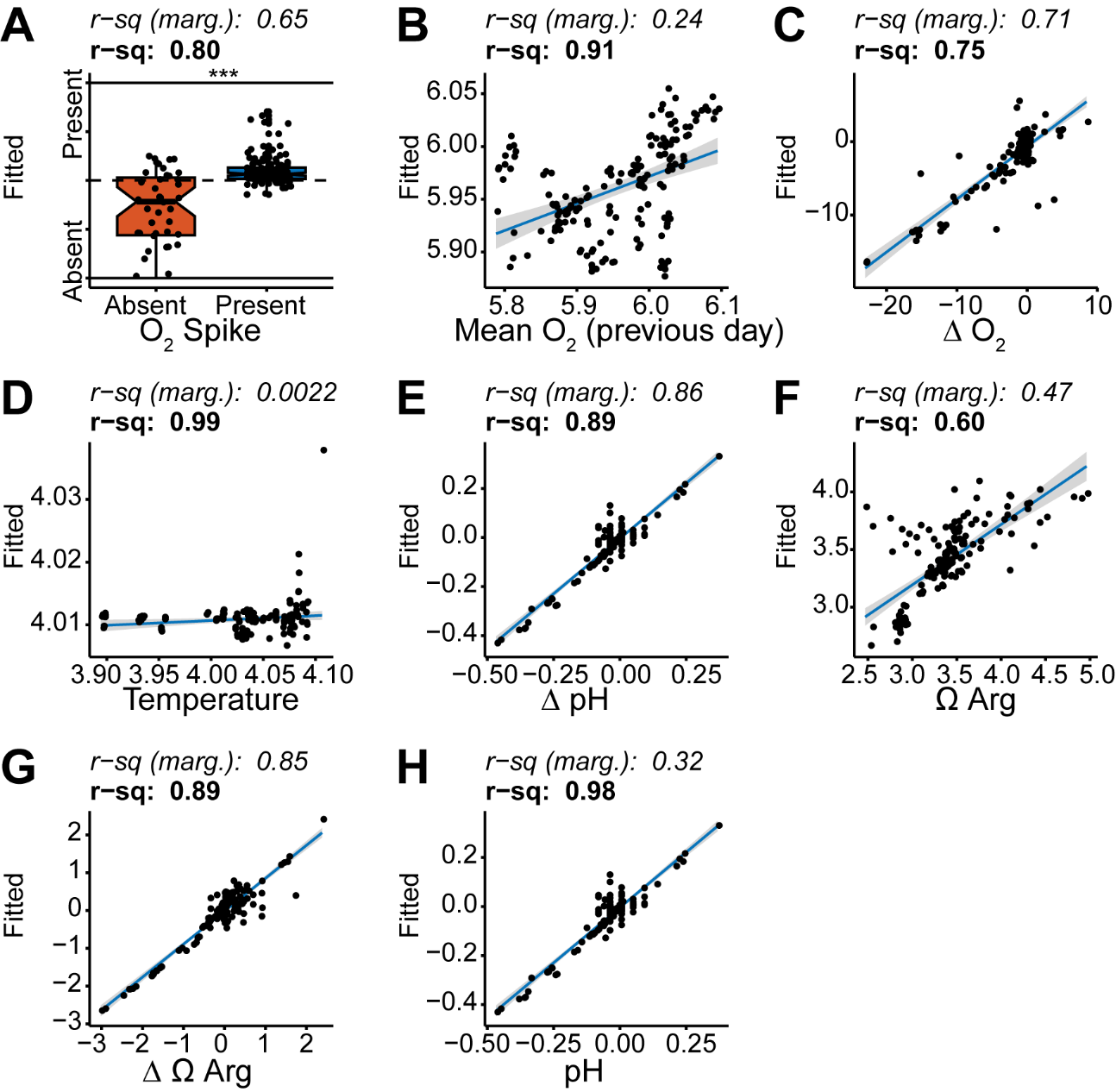

**Figure S9. Goodness of fit plots for piecewise structural equation model in Figure 3A.**

Predicted values are plotted on the y-axis and original values on the x-axis for each of the eight response variables used in the structural equation model. Blue line indicates the best fit linear regression. Gray area indicates 95% confidence interval. Asterisks indicate significantly different means as determined by a Wilcox test. Thresholds: *** = p < 0.001; ** = p < 0.01; * = p < 0.05; ns = not significant.

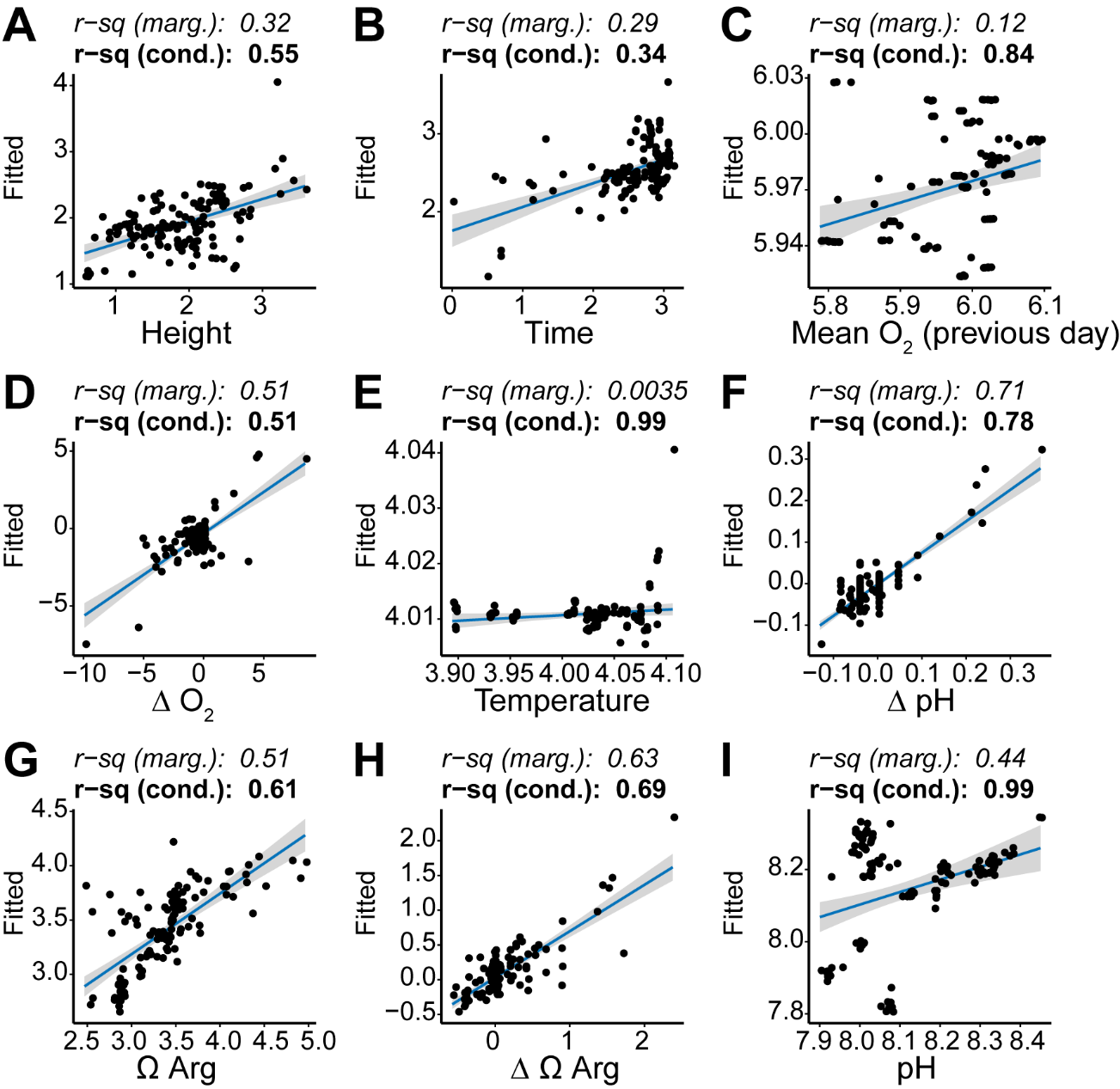

**Figure S10. Goodness of fit plots for piecewise structural equation model in Figure 3B.**

Predicted values are plotted on the y-axis and original values on the x-axis for each of the nine response variables used in the structural equation model. Blue line indicates the best fit linear regression. Gray area indicates 95% confidence interval.

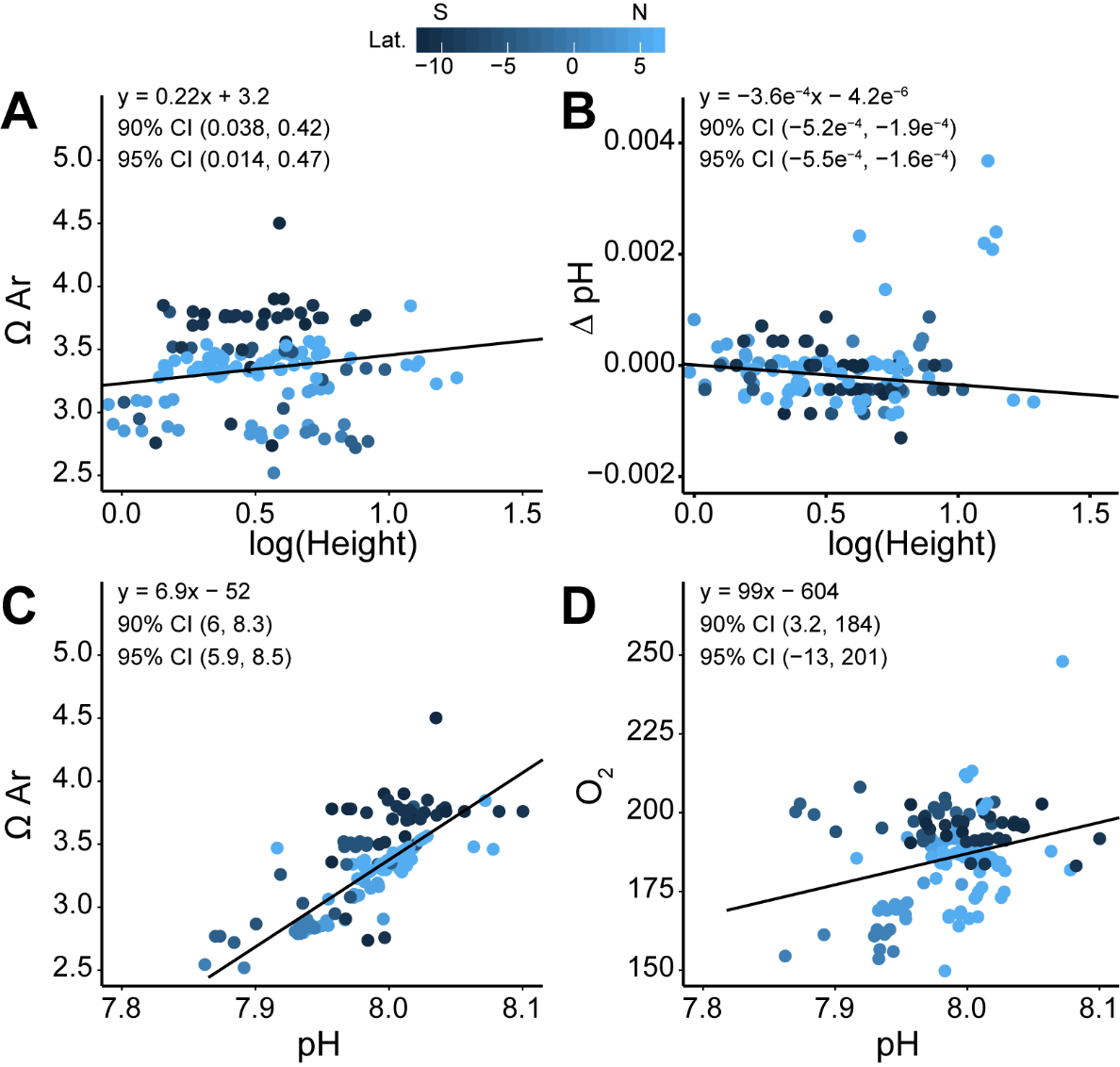

**Figure S11**. **Omega, pH and oxygen robust regressions.**

The linear formula obtained by robust regression is listed on each plot, as well as the 90% and 95% confidence intervals obtained via bootstrap of robust regression. Dotted lines indicate regressions not significant at either confidence level (a confidence interval that includes zero is not significant).

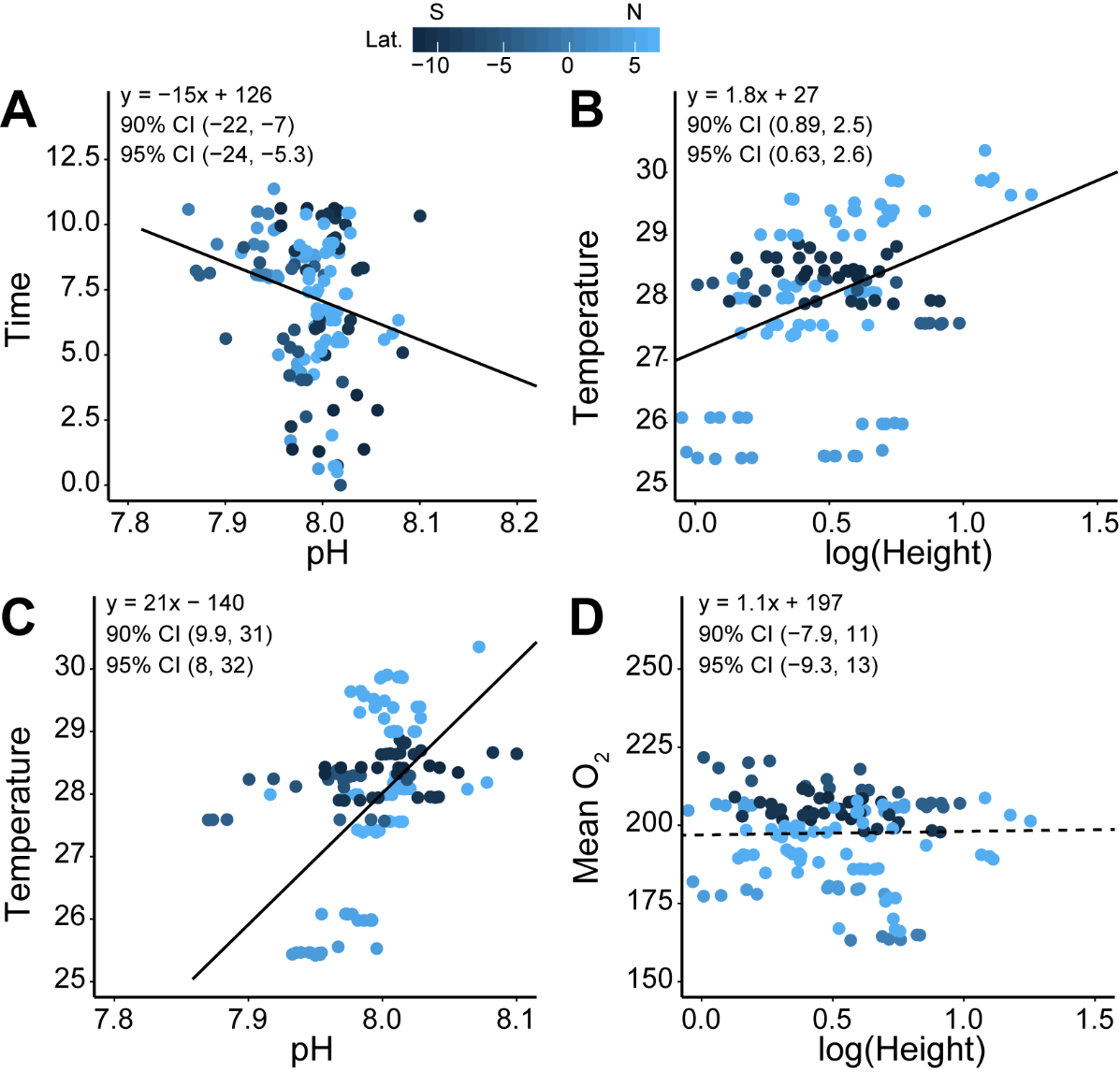

**Figure S12**. **Time and temperature robust regressions.**

The linear formula obtained by robust regression is listed on each plot, as well as the 90% and 95% confidence intervals obtained via bootstrap of robust regression. Dotted lines indicate regressions not significant at either confidence level (a confidence interval that includes zero is not significant). For panel C, no regression analysis was possible.

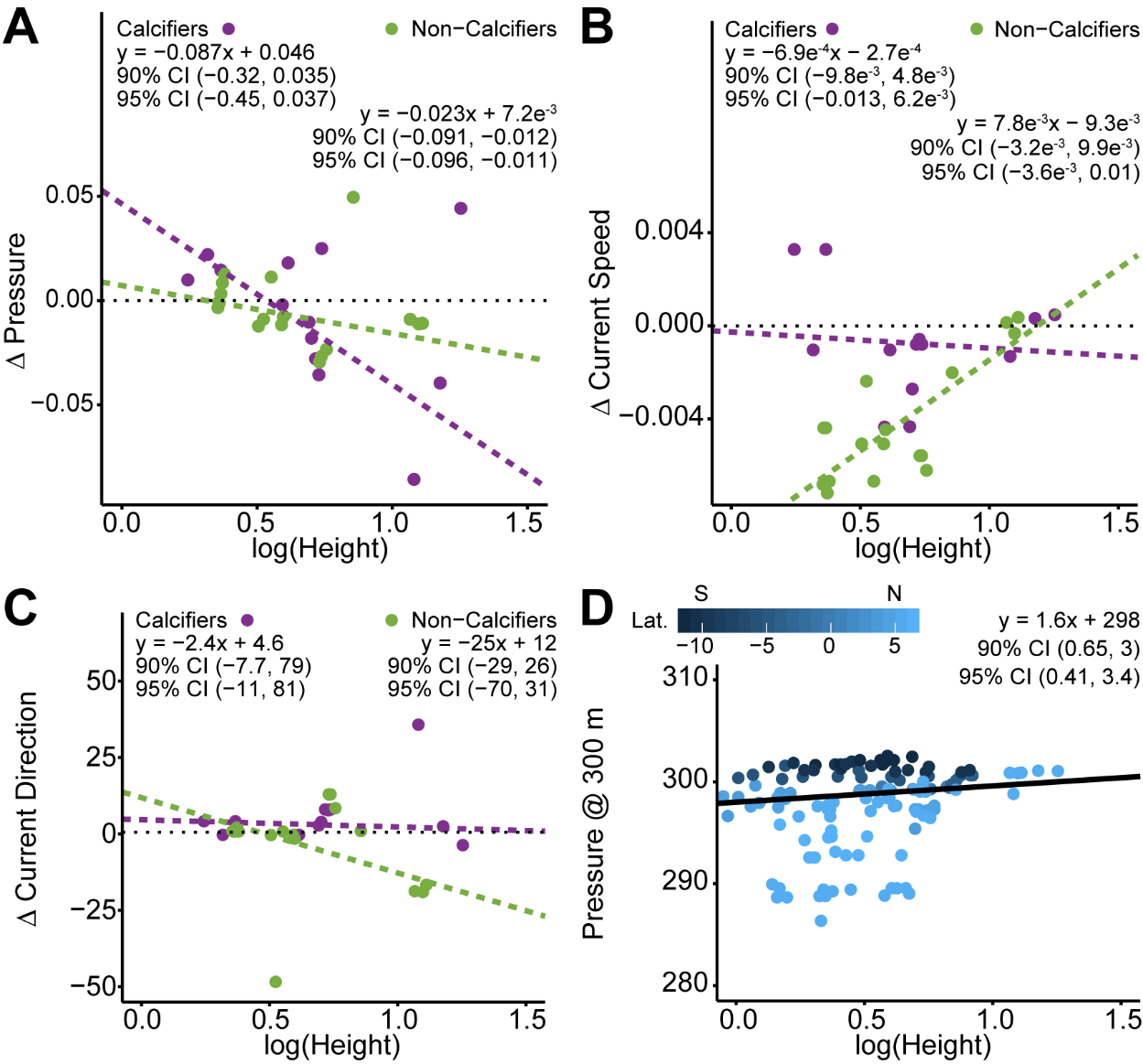

**Figure S13**. **Pressure and current robust regressions.**

The linear formula obtained by robust regression is listed on each plot, as well as the 90% and 95% confidence intervals obtained via bootstrap of robust regression. Dotted lines indicate regressions not significant at either confidence level (a confidence interval that includes zero is not significant). For panel C, no regression analysis was possible.

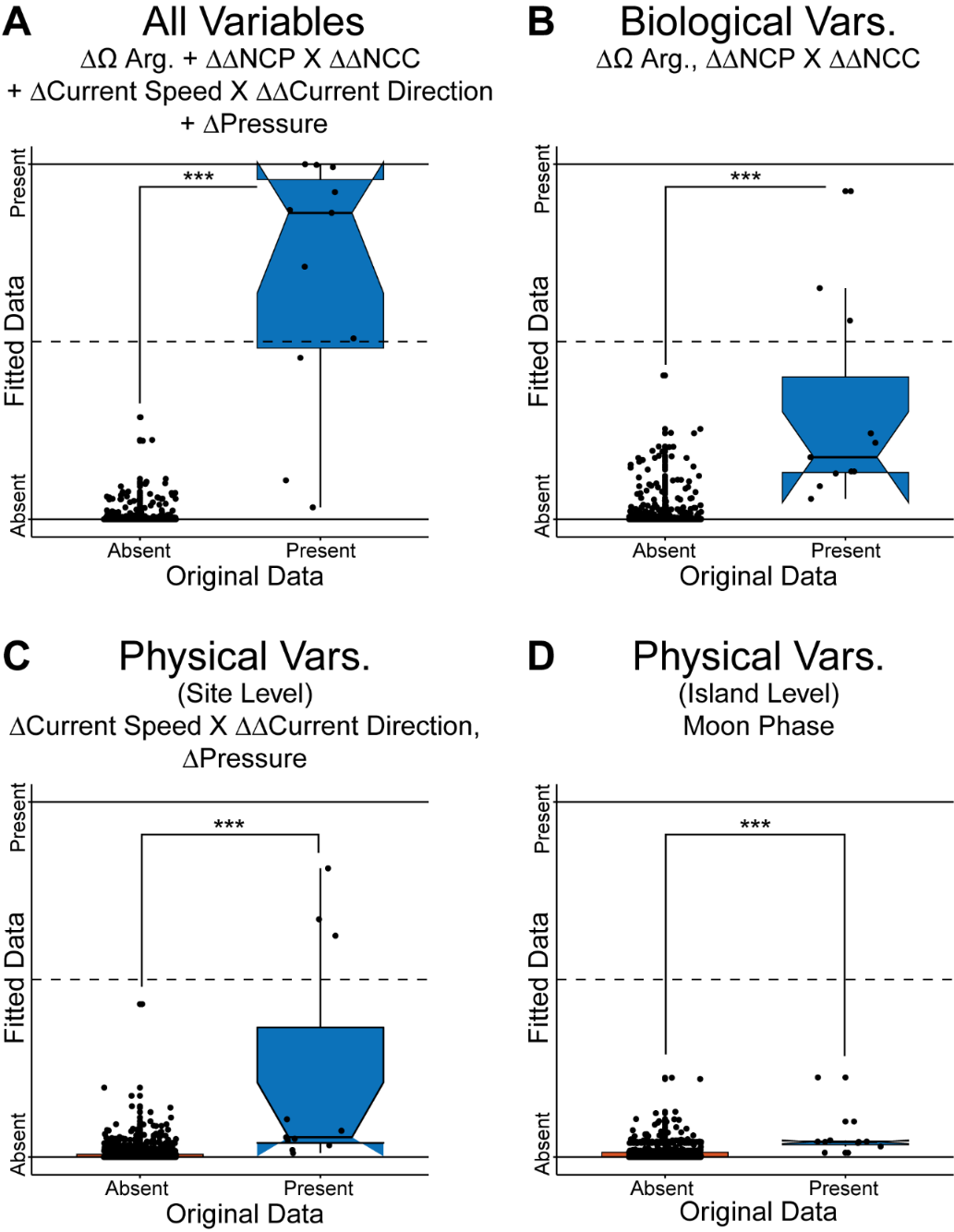

**Figure S14.** **Variables related to community production and calcification are best at predicting dissolved oxygen spikes; variables related to local currents decrease false positives.**

Best-fitting generalized additive mixed (GAM) models constructed using complete time series variables in the Palmyra BEAMS data set. Predicted values are plotted on the y-axis and original values on the x-axis. All variables were fitted as splines except where ‘x’ denotes a 2-D surface fitted to the variables on either side of the ‘x’. (A) A combination of the best fit of all variables in the other three models. Explains 81% of the variance in the data. (B) Variables predominantly influenced by biological activity. Explains 43% of the variance in the data. (C) Local, site-level variables predominantly influenced by physical forcing. Explains 40% of the variance in the data. (D) Offshore, island-level variables. Explains 41% of the variance in the data. Asterisks indicate significantly different means as determined by a Wilcox test. Thresholds: *** = p < 0.001; ** = p < 0.01; * = p < 0.05; ns = not significant.

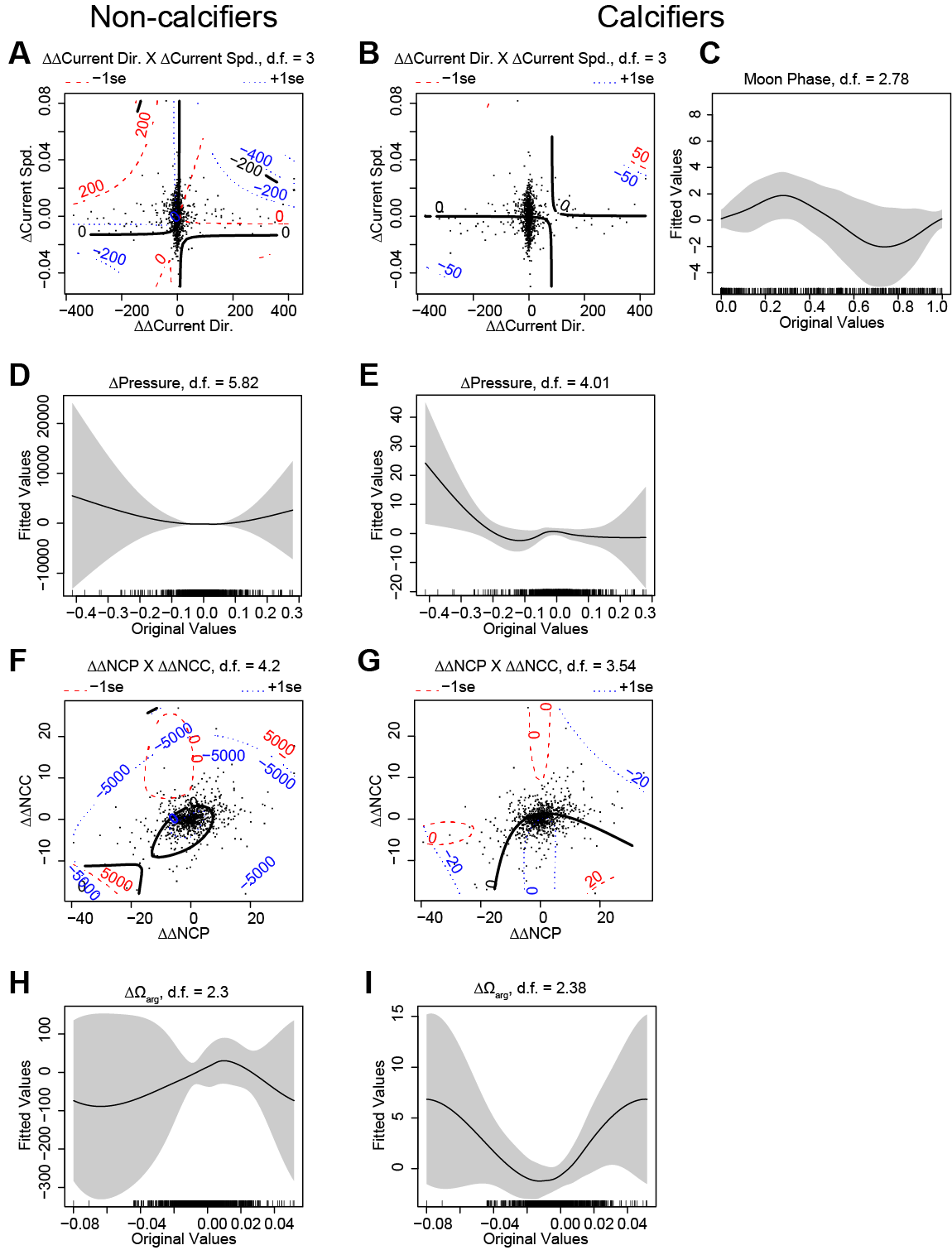

**Figure S15. Fitted splines of all GAM variables.**

Original variable values are plotted on the x-axis, and fitted spline values on the y-axis. The rug plot just above the x-axis indicates the distribution of values. The ‘s’ in front of the parenthesis signifies a 1-D spline smooth, and ‘te’ indicates a 2-D tensor (i.e. surface) smooth. The number after the variable names is the estimated degrees of freedom. Gray areas are 95% confidence intervals.

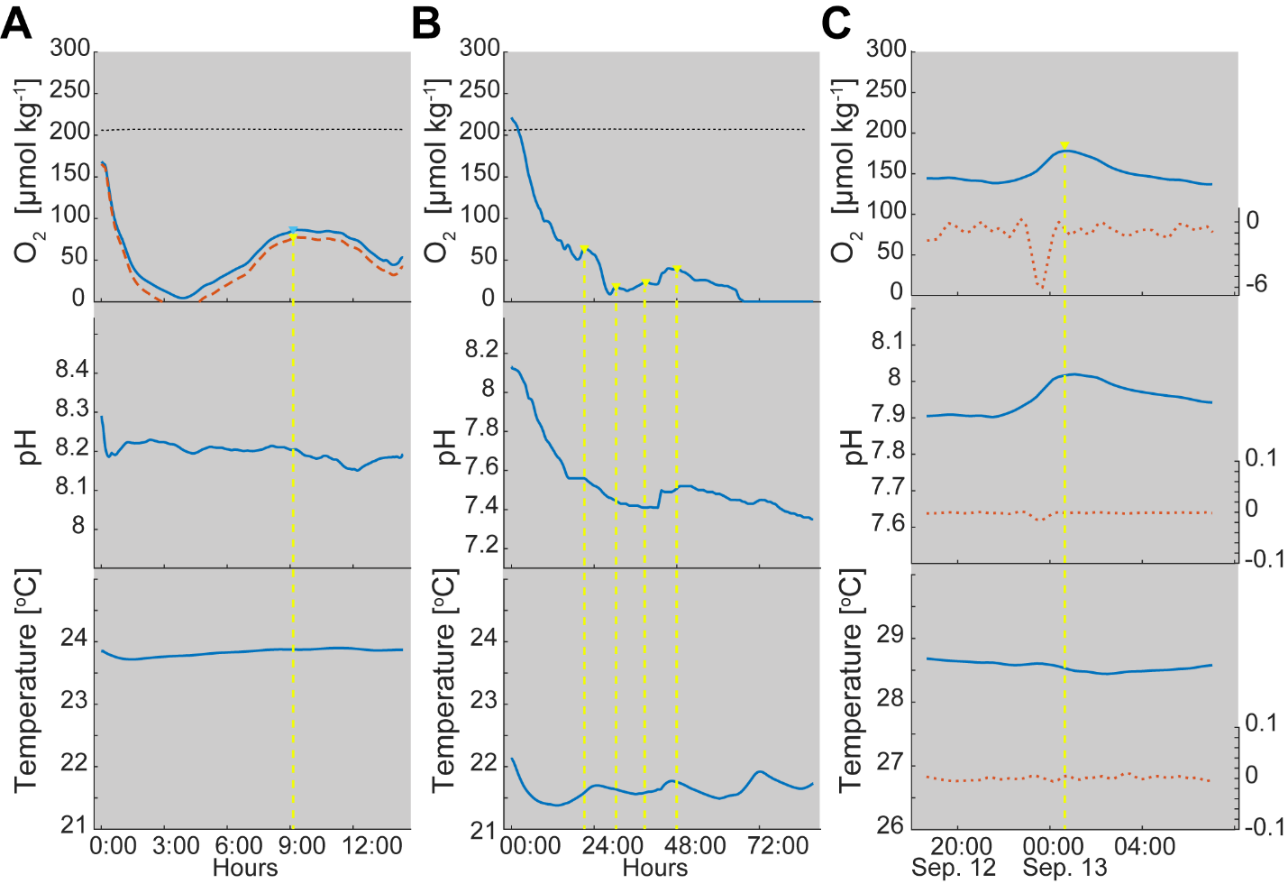

**Figure S16**. **Comparison of two of the highest oxygen spikes occurring in incubations.**

Gray background denotes darkness, yellow triangles indicate locations of oxygen spikes, and yellow dotted lines show the location of these spikes throughout the bottom graphs. Dotted black lines indicate the oxygen concentration at 100% air saturation. (A) Open incubation of CCA in CuracaoTank117. Orange dotted line shows the oxygen concentration after accounting for diffusion across the surface of the incubation chamber. Light blue triangle indicates the uncorrected oxygen spike location. (B) Closed incubation of *Montipora capricornis* coral frag in Dark Montipora Incubation.

**Supplemental Tables**

**Table S1. Location and reference data for global locations of night Time dissolved oxygen spikes in Figure 1.**

Number corresponds to the number on Figure 1. Height refers to the average Height of the nightTime oxygen spike, as estimated from the published data. Duration refers to the length of Time during which the nightTime oxygen spike occurred.

| Number | Location | Lat | Lon | Date | Height | Duration (h) | Ref. |
| --- | --- | --- | --- | --- | --- | --- | --- |
| 1 | Saca di Goro, Adriatic Sea, Italy | 44.47 N | 12.15 E | 1992 | 50% sat. | 4 to 6 | (Viaroli & Christian, 2004) |
| 2 | Long Island Sound, USA | 41.14 N | 72.77 W | 2010 | 5 µmol l^-1^ | ~6 | (Collins et al., 2013) |
| 3 | Flax Pond, Long Island Sound, USA | 40.58 N | 73.82 W | 2014 | 2 mg l^-1^ | 4 to 6 | (Baumann et al., 2014) |
| 4 | Onacock Creek, Cheasapeake Bay, USA | 37.43 N | 75.51 W | 2008 | 2 mg l^-1^ | >3 | (Shen et al., 2008) |
| 5 | South Bay, Virginia, USA | 37.15 N | 75.48 W | 2014 | 100 mmol m^-2^ d^-1^ | 4 to 6 | (Rheuban, Berg & McGlathery, 2014) |
| 6 | Eilat, Red Sea, Israel | 29.33 N | 34.57 E | 1998 | 78% sat. | ~6 | (Luz & Barkan, 2009) |
| 7 | Aqaba, Red Sea, Jordan | 29.29 N | 34.58 E | 2010 | 2 mg l^-1^ | ~6-8 | (Wild et al., 2010) |
| 8 | Florida Keys National Marine Sanctuary, USA | 25.06 N | 80.18 W | 2013 | 200 mmol m^-2^ d^-1^ | 4 | (Long et al., 2013) |
| 9 | Florida Bay, Florida, USA | 25.03 N | 80.37 W | 2000 | 200 µmol l^-1^ | 4 to 6 | (Yates et al., 2007) |
| 10 | Bora Bay, Japan | 24.45 N | 125.20 E | 1993 | 50 µmol l^-1^ | 4 | (Kraines et al., 1996) |
| 11 | Bora Bay, Japan | 24.45 N | 125.20 E | 1995 | 25 µmol l^-1^ | 4 | (Kraines et al., 1998) |
| 12 | Shiraho Reef, Ryukyu Islands, Japan | 24.22 N | 124.15 E | 1998 | 4 mg l^-1^ | ~4 | (Kayanne et al., 2008) |
| 13 | Kane'ohe Bay, O'ahu, USA | 21.25 N | 157.47 W | 2012 | 0.5 mg l^-1^ | 4 | (Martinez, Smith & Richmond, 2012) |
| 14 | Hainan, China | 19.31 N | 110.51 E | 2012 | 0.5 mg l^-1^ | ~4 | (Krumme, Herbeck & Wang, 2012) |
| 15 | Enrique Reef, La Parguera, Puerto Rico | 17.57 N | 67.03 W | 2009 | 20 µmol l^-1^ | ~6 | (McGillis et al., 2011) |
| 16 | Piscaderabai, Willemstad, Curacao | 12.7 N | 68.56 W | 2015 | 2 µmol l^-1^ | 6 | this study |
| 17 | Kingman Reef, Northern Line Islands, USA (territory) | 06.24 N | 162.24 W | 2010 | 5 µmol l^-1^ | 6 | this study |
| 18 | Palmyra Atoll, Northern Line Islands, USA (territory) | 05.52 N | 162.6 W | 2010 | 5 µmol l^-1^ | 6 | this study |
| 19 | Palmyra Atoll, Northern Line Islands, USA (territory) | 05.52 N | 162.6 W | 2014 | 15 µmol l^-1^ | 6 | this study, (Takeshita et al., 2016) |
| 20 | Teraina (Washington Island), Northern Line Islands, Kiribati | 04.43 N | 160.24 W | 2010 | 5 µmol l^-1^ | 6 | this study |
| 21 | Tabuaeran (Fanning Island), Northern Line Islands, Kiribati | 03.52 N | 159.22 W | 2010 | 5 µmol l^-1^ | 6 | this study |
| 22 | Jarvis Island, Southern Line Islands, USA (territory) | 00.22 S | 160.03 W | 2013 | 2 µmol l^-1^ | 6 | this study |
| 23 | Malden Island, Southern Line Islands, Kiribati | 04.01 S | 154.59 W | 2013 | 10 µmol l^-1^ | 6 | this study |
| 24 | Starbuck Island, Southern Line Islands, Kiribati | 05.37 S | 151.5 W | 2013 | 5 µmol l^-1^ | 6 | this study |
| 25 | Millennium Island, Southern Line Islands, Kiribati | 09.57 S | 150.13 W | 2013 | 15 µmol l^-1^ | 6 | this study |
| 26 | Vostok Island, Southern Line Islands, Kiribati | 10.06 S | 152.25 W | 2013 | 5 µmol l^-1^ | 6 | this study |
| 27 | Flint Island, Southern Line Islands, Kiribati | 11.26 S | 151.48 W | 2013 | 2 µmol l^-1^ | 6 | this study |
| 28 | Yonge Reef, Great Barrier Reef, Australia | 14.5 S | 145.6 E | 1993 | 40% sat. | 6 | (Frankignoulle et al., 1996) |
| 29 | Mo'orea, French Polynesia | 17.30 S | 145.0 W | 1975 | 1 mg l^-1^ | 4 | (Sournia, 1976) |
| 30 | Mo'orea, French Polynesia | 17.30 S | 145.0 W | 1988 | 2 mg l^-1^ | 8 | (Campion-Alsumard et al., 1993) |
| 31 | Mo'orea, French Polynesia | 17.48 S | 149.84 W | 2011 | 50 µmol l^-1^ | 6 | this study, (Haas et al., 2013) |
| 32 | Heron Island, Great Barrier Reef, Australia | 23.27 S | 151.55 E | 2011 | 50 µmol l^-1^ | 4 to 6 | (Santos et al., 2011) |

**Table S2**. **Table of all variables included from the Palmyra BEAMS and Line Islands datasets.**

The first and second derivatives for all time series variables were also used.

|  |  | Palmyra BEAMS | | Line Islands cBITs | |
| --- | --- | --- | --- | --- | --- |
| Variable | Specificity | Collection Freq. | Collection Method | Collection Freq. | Collection Method |
| Dissolved Oxygen | Site | 12 min. | SeapHOx Multi-sensor Sonde | 5 min. | MANTA Multi-sensor Sonde |
| pH | Site | 12 min. | SeapHOx Multi-sensor Sonde | 5 min. | MANTA Multi-sensor Sonde |
| Temperature | Site | 12 min. | SeapHOx Multi-sensor Sonde | 5 min. | MANTA Multi-sensor Sonde |
| ORP | Site | NA | NA | 5 min. | MANTA Multi-sensor Sonde |
| Conductivity | Site | 12 min. | SeapHOx Multi-sensor Sonde | 5 min. | MANTA Multi-sensor Sonde |
| Ω Aragonite | Site | 12 min. | Calculated from discreet and Time series data | Daily | Calculated from discreet samples |
| Net Community Calcification | Site | 12 min. | Calculated from discreet and Time series data | NA | NA |
| Net Community Production | Site | 12 min. | Calculated from discreet and Time series data | NA | NA |
| Current Speed | Site | 12 min. | Acoustic Doppler Velocimeter | NA | NA |
| Current Direction | Site | 12 min. | Acoustic Doppler Velocimeter | NA | NA |
| Pressure | Site | 12 min. | Acoustic Doppler Velocimeter | NA | NA |
| Photosynthetically Active Radation | Island | 12 min. | LICOR 4-pi Sensor | 1 min | LICOR 4-pi Sensor |
| Moon Phase | Island | Hourly | U.S. Naval Charts | Hourly | U.S. Naval Charts |
| Offshore Pressure @ 300 m | Island | Hourly | NOAA Buoy | Hourly | NOAA Buoy |
| Offshore Pressure @ 500 m | Island | Hourly | NOAA Buoy | Hourly | NOAA Buoy |
| Wind Speed | Island | Hourly | NOAA Buoy | Hourly | NOAA Buoy |
| Wind Direction | Island | Hourly | NOAA Buoy | Hourly | NOAA Buoy |
| Lattitude | Island | Once | GPS (site level) | Once | GPS |
| Longitude | Island | Once | GPS (site level) | Once | GPS |
| Crustose Coralline Algae | Site | Once | Photoquadrat | Once | Photoquadrat |
| Macroalgae | Site | Once | Photoquadrat | Once | Photoquadrat |
| Hallimeda | Site | NA | NA | Once | Photoquadrat |
| Mixed Turf Algae | Site | NA | Photoquadrat | Once | Photoquadrat |
| Hard Coral | Site | Once | Photoquadrat | Once | Photoquadrat |
| Soft Coral | Site | NA | NA | Once | Photoquadrat |
| Corallimorph | Site | Once | Photoquadrat | NA | NA |
| Other Calcifiers | Site | Once | Photoquadrat | Once | Photoquadrat |
| Peyssonnelia | Site | NA | NA | Once | Photoquadrat |
| Black Crust | Site | NA | NA | Once | Photoquadrat |
| Tridacna | Site | NA | NA | Once | Photoquadrat |
| Non-biological | Site | NA | NA | Once | Photoquadrat |

**Table S3. Summary of oxygen spike detection algorithm for all datasets.**

| Dataset | Present | Absent | Total |
| --- | --- | --- | --- |
| Curacao | 4 | 2 | 6 |
| Mo'orea | 41 | 13 | 54 |
| Line Islands cBITs | 142 | 20 | 162 |
| Palmyra BEAMS | 44 | 18 | 62 |
| Total | 231 | 53 | 284 |
| LI + Palmyra | 186 | 38 | 224 |

**Table S4. Summary statistics for oxygen spikes across all datasets.**

|  | Oxygen [µmol kg^-1^] | Height [µmol kg^-1^] | Width [h] |
| --- | --- | --- | --- |
| Mean | 176.9 +/- 24.3 | 5.1 +/- 5.2 | 2.6 +/- 1.2 |
| Median | 184.7 | 3.8 | 2.4 |
| Max | 243.6 | 35.1 | 8.0 |
| Dataset (max) | Palmyra | Mo'orea | Line Islands |
| Time (max) | 9/15/2014 0:55 | 9/9/2011 1:15 | 10/24/2013 2:30 |
| **Curacao** |  |  |  |
| Mean | 176.9 +/- 0.9 | 1.5 +/- 1.0 | 1.9 +/- 0.4 |
| Median | 176.8 | 1.0 | 1.7 |
| Max | 178.5 | 3.5 | 2.8 |
| Time (max) | 5/9/2015 22:10 | 5/12/2015 2:30 | 5/12/2015 2:30 |
| **Mo'orea** |  |  |  |
| Mean | 137.0 +/- 20.2 | 8.6 +/- 7.4 | 3.0 +/- 1.3 |
| Median | 139.0 | 5.6 | 2.8 |
| Time (max) | 183.1 | 35.1 | 5.8 |
| Time | 9/5/2011 19:39 | 9/9/2011 1:15 | 9/6/2011 2:40 |
| **Line Islands** |  |  |  |
| Mean | 188.0 +/- 11.4 | 3.6 +/- 2.1 | 2.6 +/- 1.2 |
| Median | 190.7 | 3.3 | 2.4 |
| Max | 208.1 | 10.8 | 8.0 |
| Time (max) | 10/29/2013 3:35 | 11/6/2013 2:35 | 10/24/2013 2:30 |
| **Palmyra** |  |  |  |
| Mean | 178.2 +/- 20.7 | 7.3 +/- 7.1 | 2.5 +/- 1.2 |
| Median | 173.1 | 5.1 | 2.2 |
| Max | 243.6 | 34.0 | 6.1 |
| Time (max) | 9/15/2014 0:55 | 9/13/2014 1:04 | 9/12/2014 23:37 |

**Table S5. Models used for structural equation model in Figure 3A.**

Island was used as a random variable in each.

| Method | Response | Predictors | AIC | AICc | df | p-value |
| --- | --- | --- | --- | --- | --- | --- |
| glmer, binary | O2 Spike +/- | Δ Ω Ar + ΔΔ pH + O2 | -- | -- | -- | -- |
| lme, gaussian | O2 | TEMP. + ΔΔ TEMP. + Δ Ω Ar + Mean O2 + O2:PAR + ΔΔ O2 | -- | -- | -- | -- |
| lme, gaussian | Δ O2 | TEMP. + ΔΔ TEMP. + Δ pH + ΔΔ pH + Mean O2 + ΔΔ O2 | -- | -- | -- | -- |
| lme, gaussian | TEMP. | ΔΔ TEMP. + Mean PAR | -- | -- | -- | -- |
| lme, gaussian | Δ pH | Δ O2 + ΔΔ O2 + Δ Ω Ar + ΔΔ pH + ΔΔ TEMP. | -- | -- | -- | -- |
| lme, gaussian | Ω Ar | pH + O2:PAR + Mean PAR + TEMP. | -- | -- | -- | -- |
| lme, gaussian | Δ Ω Ar | Δ pH + ΔΔ pH + O2 + Δ O2 + ΔΔ O2 + Mean O2 | -- | -- | -- | -- |
| -- | -- | -- | 233.07 | 283.77 | 80 | 0.004 |

**Table S6. SEM coefficients for structural equation model in Figure 3A.**

| Response | Predictor | Coefficient | Std. Error | p-value |
| --- | --- | --- | --- | --- |
| O2 Spike +/- | O2 | 33.64 | 10.3752 | 0.0012 |
| Δ O2 | Δ pH | 24.75 | 2.11356 | 0 |
| Δ O2 | Mean O2 | 10.12 | 3.89625 | 0.0102 |
| O2 Spike +/- | Δ Ω Ar | 9.92 | 3.16853 | 0.0017 |
| Ω Ar | pH | 8.58 | 0.90276 | 0 |
| Δ Ω Ar | Δ pH | 5.25 | 0.26673 | 0 |
| Δ O2 | ΔΔ Temp. | 4.97 | 0.6192 | 0 |
| O2 | Temp. | 1.52 | 0.30825 | 0 |
| Δ Ω Ar | ΔΔ pH | 1.34 | 0.46602 | 0.0044 |
| Δ O2 | Temp. | -22.67 | 6.75281 | 0.001 |
| O2 Spike +/- | ΔΔ pH | -33.28 | 11.3048 | 0.0032 |

**Table S7. Models used for structural equation model in Figure 3B.**

Island was used as a random variable in each model.

| Method | Response | Predictors | AIC | AICc | Df | p-value |
| --- | --- | --- | --- | --- | --- | --- |
| lme, gaussian | Height | ΔΔ Ω Ar + O2:PAR + Mean PAR + Δ O2 + Mean O2 + Time + Δ pH + Δ Temp. | -- | -- | -- | -- |
| lme, gaussian | Time | O2 + Δ O2 + Height + pH + Mean PAR | -- | -- | -- | -- |
| lme, gaussian | O2 | Temp. + Δ Temp. + Mean O2 + O2:PAR | -- | -- | -- | -- |
| lme, gaussian | Δ Temp. | O2 + O2:PAR + Mean O2 + Temp. + ΔΔ Temp. | -- | -- | -- | -- |
| lme, gaussian | Δ pH | O2 + O2:PAR + Δ Ω + ΔΔ Ω Ar + ΔΔ pH + Δ Temp. + ΔΔ Temp. | -- | -- | -- | -- |
| lme, gaussian | Ω Ar | pH + ΔΔ Ω Ar + O2 + Mean PAR + Temp. | -- | -- | -- | -- |
| lme, gaussian | Δ Ω Ar | Ω Ar + ΔΔ Ω Ar + ΔΔ Temp. + Δ pH + ΔΔ pH + O2 + Δ O2 + O2:PAR | -- | -- | -- | -- |
| lme, gaussian | pH | Δ Ω Ar + Ω Ar | -- | -- | -- | -- |
| -- | -- | -- | 256.1 | 416.43 | 104 | 0.409 |

**Table S8. SEM coefficients for structural equation model in Figure 3B.**

| Response | Predictor | Coefficient | Std. Error | p-value |
| --- | --- | --- | --- | --- |
| Height | Mean O2 | 13.33 | 4.29E+00 | 0.0023 |
| Δ Temp. | O2 | 9.12 | 6.62E-01 | 0 |
| Ω Ar | pH | 8.59 | 1.16E+00 | 0 |
| Δ Ω Ar | Δ pH | 3.74 | 3.89E-01 | 0 |
| Δ Temp. | Mean O2 | 2.89 | 8.02E-01 | 0.0004 |
| Height | Δ Temp. | 2.57 | 3.82E-01 | 0 |
| Δ Ω Ar | O2 | 1.57 | 4.02E-01 | 0.0002 |
| Δ Ω Ar | ΔΔ pH | 1.39 | 4.71E-01 | 0.0039 |
| O2 | Temp. | 1.36 | 2.51E-01 | 0 |
| Time | Mean PAR | -1.05 | 4.11E-01 | 0.0115 |
| Height | Mean PAR | -2.67 | 3.62E-01 | 0 |
| Height | O2:PAR | -2.83 | 4.69E-01 | 0 |
| Height | ΔΔ Ω Ar | -6.94 | 1.41E+00 | 0 |
| Δ Temp. | Temp. | -7.29 | 2.90E+00 | 0.0133 |
| Time | O2 | -11.79 | 2.79E+00 | 0 |
| Height | Δ pH | -15.39 | 3.75E+00 | 0.0001 |
| Time | pH | -124.84 | 4.47E+01 | 0.006 |

**Table S9. Oxygen spike positive incubation details.**

| Incubation | Organism | Vol_ratio | Source | Light | Orientation | Sealed | Sensor_pos | Organism_pos |
| --- | --- | --- | --- | --- | --- | --- | --- | --- |
| CuracaoTank316 | CCA | 0.25 | Field collection, 12 m | Darkness | Vertical | N | Top | Bottom |
| CuracaoTank117 | CCA | 0.25 | Field collection, intertidal | Darkness | Vertical | N | Top | Bottom |
| Dark Montipora Incubation | Montipora capricornis | 0.1 | Aquaculture | Darkness | Horizontal | Y | Side | MiΔΔ le |
| Montipora Incubation 1 | Montipora capricornis | 0.1 | Aquaculture | Diurnal cycle | Vertical | Y | Top | Bottom |
| Dark Montipora Incubation | Montipora capricornis | 0.1 | Aquaculture | Darkness | Horizontal | Y | Side | MiΔΔ le |
| Montipora Incubation 3 | Montipora capricornis | 0.1 | Aquaculture | Diurnal cycle | Horizontal | Y | Side | MiΔΔ le |
| Montipora Incubation 2 | Montipora capricornis | 0.1 | Aquaculture | Diurnal cycle | Vertical | Y | Top | Bottom |
| Dark Montipora Incubation | Montipora capricornis | 0.1 | Aquaculture | Darkness | Horizontal | Y | Side | MiΔΔ le |
| Dark Montipora Incubation | Montipora capricornis | 0.1 | Aquaculture | Darkness | Horizontal | Y | Side | MiΔΔ le |
| Montipora Incubation 1 | Montipora capricornis | 0.1 | Aquaculture | Diurnal cycle | Vertical | Y | Top | Bottom |
| Montipora Incubation 3 | Montipora capricornis | 0.1 | Aquaculture | Diurnal cycle | Horizontal | Y | Side | MiΔΔ le |
| CuracaoEpilith3 | CCA | 0.01 | Field collection, intertidal | Diurnal cycle | Vertical | Y | Top | Bottom |
| Montipora Incubation 2 | Montipora capricornis | 0.1 | Aquaculture | Diurnal cycle | Vertical | Y | Top | Bottom |
| CuracaoEndolith3 | CCA | 0.01 | Field collection, intertidal | Diurnal cycle | Vertical | Y | Bottom | Bottom |
| CuracaoEpilith2 | CCA | 0.01 | Field collection, intertidal | Diurnal cycle | Vertical | Y | Top | Bottom |
| Montipora Incubation 1 | Montipora capricornis | 0.1 | Aquaculture | Diurnal cycle | Vertical | Y | Top | Bottom |
| CuracaoEpilith1 | CCA | 0.01 | Field collection, intertidal | Diurnal cycle | Vertical | Y | Top | Bottom |
| CuracaoEpilith3 | CCA | 0.01 | Field collection, intertidal | Diurnal cycle | Vertical | Y | Top | Bottom |
| CuracaoEpilith4 | CCA | 0.01 | Field collection, intertidal | Diurnal cycle | Vertical | Y | Bottom | Bottom |
| CuracaoEpilith1 | CCA | 0.01 | Field collection, intertidal | Diurnal cycle | Vertical | Y | Top | Bottom |
| CuracaoEpilith4 | CCA | 0.01 | Field collection, intertidal | Diurnal cycle | Vertical | Y | Bottom | Bottom |
| CuracaoEndolith2 | CCA | 0.01 | Field collection, intertidal | Diurnal cycle | Vertical | Y | Bottom | Bottom |
| Montipora Incubation 1 | Montipora capricornis | 0.1 | Aquaculture | Diurnal cycle | Vertical | Y | Top | Bottom |
| CuracaoTank314 | Seawater | 1 | Field collection, 0 m | Darkness | Vertical | N | Top | Bottom |
| CuracaoEpilith0 | CCA | 0.01 | Field collection, intertidal | Diurnal cycle | Vertical | Y | Top | Bottom |
| CuracaoEndolith0 | CCA | 0.01 | Field collection, intertidal | Diurnal cycle | Vertical | Y | Top | Bottom |
| CuracaoEndolith0 | CCA | 0.01 | Field collection, intertidal | Diurnal cycle | Vertical | Y | Top | Bottom |
| CuracaoEndolith1 | CCA | 0.01 | Field collection, intertidal | Diurnal cycle | Vertical | Y | Bottom | Bottom |
| CuracaoEndolith1 | CCA | 0.01 | Field collection, intertidal | Diurnal cycle | Vertical | Y | Bottom | Bottom |
| CuracaoTank44 | RO water | 1 | Tap | Darkness | Vertical | N | Top | Bottom |
| CuracaoTank44 | RO water | 1 | Tap | Darkness | Vertical | N | Top | Bottom |
| CuracaoTank29 | Mixed | 0.67 | Field collection, 10 m | Darkness | Vertical | N | Top | Bottom |
| CuracaoTank19 | Mixed | 0.67 | Field collection, 10 m | Darkness | Vertical | N | Top | Bottom |
| CuracaoTank212 | Turf | 0.5 | Field collection, 12 m | Darkness | Vertical | N | Top | Bottom |
| CuracaoTank19 | Mixed | 0.67 | Field collection, 10 m | Darkness | Vertical | N | Top | Bottom |

**Table S10. Oxygen spike information for all positive incubations.**

| Incubation | Organism | O2 | Height | Adj_Height | Width | Time |
| --- | --- | --- | --- | --- | --- | --- |
| CuracaoTank316 | CCA | 76.8 | 46.9 | 37.2 | 2.1 | 7.4 |
| CuracaoTank117 | CCA | 87.2 | 43.3 | 34.3 | 5 | 9.1 |
| Dark Montipora Incubation | Montipora capricornis | 40.4 | 31.2 | 31.2 | 14.5 | 48.1 |
| Montipora Incubation 1 | Montipora capricornis | 490.7 | 17.3 | 17.3 | 1.8 | 312.1 |
| Dark Montipora Incubation | Montipora capricornis | 64.6 | 13.8 | 13.8 | 3.3 | 21.2 |
| Montipora Incubation 3 | Montipora capricornis | 307.1 | 10 | 10 | 2.6 | 27.9 |
| Montipora Incubation 2 | Montipora capricornis | 209.7 | 9.1 | 9.1 | 1.3 | 44.9 |
| Dark Montipora Incubation | Montipora capricornis | 19 | 5.9 | 5.9 | 1.8 | 30.4 |
| Dark Montipora Incubation | Montipora capricornis | 24 | 3.1 | 3.1 | 1.9 | 39 |
| Montipora Incubation 1 | Montipora capricornis | 211.4 | 3 | 3 | 2.5 | 2.3 |
| Montipora Incubation 3 | Montipora capricornis | 375.9 | 2.8 | 2.8 | 1.1 | 48 |
| CuracaoEpilith3 | CCA | 117 | 2.7 | 2.7 | 2.3 | 24 |
| Montipora Incubation 2 | Montipora capricornis | 213.6 | 2.6 | 2.6 | 4.8 | 1.7 |
| CuracaoEndolith3 | CCA | 49 | 2.4 | 2.4 | 1.1 | 240.1 |
| CuracaoEpilith2 | CCA | 123.9 | 2.4 | 2.4 | 1.5 | 44 |
| Montipora Incubation 1 | Montipora capricornis | 358.5 | 2.4 | 2.4 | 2 | 53.3 |
| CuracaoEpilith1 | CCA | 152.2 | 2.2 | 2.2 | 6.2 | 30.8 |
| CuracaoEpilith3 | CCA | 110.6 | 2.1 | 2.1 | 1.8 | 47.5 |
| CuracaoEpilith4 | CCA | 102.1 | 1.9 | 1.9 | 2.7 | 13.9 |
| CuracaoEpilith1 | CCA | 144.2 | 1.7 | 1.7 | 1.7 | 53 |
| CuracaoEpilith4 | CCA | 102.1 | 1.6 | 1.6 | 2.5 | 39.2 |
| CuracaoEndolith2 | CCA | 87.9 | 1.5 | 1.5 | 3.1 | 38.3 |
| Montipora Incubation 1 | Montipora capricornis | 223.1 | 1.5 | 1.5 | 1.9 | 54.1 |
| CuracaoTank314 | Seawater | 165.1 | 2.7 | 1.4 | 1.7 | 0.9 |
| CuracaoEpilith0 | CCA | 92.8 | 1.1 | 1.1 | 1.2 | 30.9 |
| CuracaoEndolith0 | CCA | 135.5 | 0.9 | 0.9 | 1.2 | 29.7 |
| CuracaoEndolith0 | CCA | 120.4 | 0.7 | 0.7 | 1.6 | 6.1 |
| CuracaoEndolith1 | CCA | 98.7 | 0.6 | 0.6 | 1.4 | 41.4 |
| CuracaoEndolith1 | CCA | 165 | 0.6 | 0.6 | 1.2 | 3.5 |
| CuracaoTank44 | RO water | 204.7 | 0.7 | 0 | 3.3 | 49.8 |
| CuracaoTank44 | RO water | 205 | 0.8 | 0 | 2.5 | 11.6 |
| CuracaoTank29 | Mixed | 27.6 | 6.1 | 0 | 4.1 | 11.3 |
| CuracaoTank19 | Mixed | 18.8 | 9.5 | 0 | 1 | 13.1 |
| CuracaoTank212 | Turf | 52.2 | 9.7 | 0 | 1.4 | 6.3 |
| CuracaoTank19 | Mixed | 11.7 | 10.3 | 0 | 1.2 | 15.9 |

**Table S11. Oxygen spike negative incubation details.**

| Incubation | Organism | Vol_ratio | Source | Light | Orientation | Sealed | Sensor_pos | Organism_pos |
| --- | --- | --- | --- | --- | --- | --- | --- | --- |
| CuracaoCCA | CCA | 0.1 | Field collection, intertidal | Diurnal cycle | Vertical | Y | Top | Bottom |
| CuracaoCCA | CCA | 0.1 | Field collection, intertidal | Diurnal cycle | Vertical | Y | Top | Bottom |
| CuracaoCCA | CCA | 0.1 | Field collection, intertidal | Diurnal cycle | Vertical | Y | Top | Bottom |
| CuracaoEndolith0 | CCA | 0.01 | Field collection, intertidal | Diurnal cycle | Vertical | Y | Top | Bottom |
| CuracaoEndolith1 | CCA | 0.01 | Field collection, intertidal | Diurnal cycle | Vertical | Y | Bottom | Bottom |
| CuracaoEndolith1 | CCA | 0.01 | Field collection, intertidal | Diurnal cycle | Vertical | Y | Bottom | Bottom |
| CuracaoEndolith2 | CCA | 0.01 | Field collection, intertidal | Diurnal cycle | Vertical | Y | Bottom | Bottom |
| CuracaoEndolith2 | CCA | 0.01 | Field collection, intertidal | Diurnal cycle | Vertical | Y | Bottom | Bottom |
| CuracaoEndolith3 | CCA | 0.01 | Field collection, intertidal | Diurnal cycle | Vertical | Y | Bottom | Bottom |
| CuracaoEndolith3 | CCA | 0.01 | Field collection, intertidal | Diurnal cycle | Vertical | Y | Bottom | Bottom |
| CuracaoEndolith3 | CCA | 0.01 | Field collection, intertidal | Diurnal cycle | Vertical | Y | Bottom | Bottom |
| CuracaoEpilith0 | CCA | 0.01 | Field collection, intertidal | Diurnal cycle | Vertical | Y | Top | Bottom |
| CuracaoEpilith0 | CCA | 0.01 | Field collection, intertidal | Diurnal cycle | Vertical | Y | Top | Bottom |
| CuracaoEpilith1 | CCA | 0.01 | Field collection, intertidal | Diurnal cycle | Vertical | Y | Top | Bottom |
| CuracaoEpilith2 | CCA | 0.01 | Field collection, intertidal | Diurnal cycle | Vertical | Y | Top | Bottom |
| CuracaoEpilith2 | CCA | 0.01 | Field collection, intertidal | Diurnal cycle | Vertical | Y | Top | Bottom |
| CuracaoEpilith3 | CCA | 0.01 | Field collection, intertidal | Diurnal cycle | Vertical | Y | Top | Bottom |
| CuracaoEpilith4 | CCA | 0.01 | Field collection, intertidal | Diurnal cycle | Vertical | Y | Bottom | Bottom |
| CuracaoTank116 | CCA | 0.1 | Field collection, 12 m | Darkness | Vertical | N | Top | Bottom |
| CuracaoTank216 | CCA | 0.1 | Field collection, 12 m | Darkness | Vertical | N | Top | Bottom |
| CuracaoTank217 | CCA | 0.25 | Field collection, Intertidal | Darkness | Vertical | N | Top | Bottom |
| CuracaoTank317 | CCA | 0.25 | Field collection, Intertidal | Darkness | Vertical | N | Top | Bottom |
| CuracaoTank318 | CCA | 0.25 | Field collection, 12 m | Darkness | Vertical | N | Top | Bottom |
| CuracaoTank413 | CCA | 0.1 | Field collection, 5 m | Darkness | Vertical | N | Top | Bottom |
| CuracaoTank416 | CCA | 0.1 | Field collection, 12 m | Darkness | Vertical | N | Top | Bottom |
| CuracaoTank418 | CCA | 0.25 | Field collection, 12 m | Darkness | Vertical | N | Top | Bottom |
| CuracaoTank519 | CCA | 0.25 | Field collection, 12 m | Darkness | Vertical | N | Top | Bottom |
| CuracaoTank513 | Dry rubble | 0.5 | Field collection, intertidal | Darkness | Vertical | N | Top | Bottom |
| CuracaoTank34 | Seawater | 1 | Field collection, 9 m | Darkness | Vertical | N | Top | Bottom |
| CuracaoTank15 | Mixed | 0.67 | Field collection, 9 m | Darkness | Vertical | N | Top | Bottom |
| CuracaoTank17 | Mixed | 0.5 | Field collection, 10 m | Darkness | Vertical | N | Top | Bottom |
| CuracaoTank27 | Mixed | 0.5 | Field collection, 10 m | Darkness | Vertical | N | Top | Bottom |
| CuracaoTank28 | Mixed | 0.67 | Field collection, 10 m | Darkness | Vertical | N | Top | Bottom |
| CuracaoTank37 | Mixed | 0.5 | Field collection, 10 m | Darkness | Vertical | N | Top | Bottom |
| CuracaoTank38 | Mixed | 0.67 | Field collection, 10 m | Darkness | Vertical | N | Top | Bottom |
| CuracaoTank39 | Mixed | 0.67 | Field collection, 10 m | Darkness | Vertical | N | Top | Bottom |
| CuracaoTank49 | Mixed | 0.67 | Field collection, 10 m | Darkness | Vertical | N | Top | Bottom |
| CuracaoTank59 | Mixed | 0.67 | Field collection, 10 m | Darkness | Vertical | N | Top | Bottom |
| Montipora Incubation 1 | Montipora capricornis | 0.1 | Aquaculture | Diurnal cycle | Vertical | Y | Top | Bottom |
| Montipora Incubation 1 | Montipora capricornis | 0.1 | Aquaculture | Diurnal cycle | Vertical | Y | Top | Bottom |
| Montipora Incubation 1 | Montipora capricornis | 0.1 | Aquaculture | Diurnal cycle | Vertical | Y | Top | Bottom |
| Montipora Incubation 1 | Montipora capricornis | 0.1 | Aquaculture | Diurnal cycle | Vertical | Y | Top | Bottom |
| Montipora Incubation 1 | Montipora capricornis | 0.1 | Aquaculture | Diurnal cycle | Vertical | Y | Top | Bottom |
| Montipora Incubation 1 | Montipora capricornis | 0.1 | Aquaculture | Diurnal cycle | Vertical | Y | Top | Bottom |
| Montipora Incubation 1 | Montipora capricornis | 0.1 | Aquaculture | Diurnal cycle | Vertical | Y | Top | Bottom |
| Montipora Incubation 1 | Montipora capricornis | 0.1 | Aquaculture | Diurnal cycle | Vertical | Y | Top | Bottom |
| Montipora Incubation 1 | Montipora capricornis | 0.1 | Aquaculture | Diurnal cycle | Vertical | Y | Top | Bottom |
| Montipora Incubation 1 | Montipora capricornis | 0.1 | Aquaculture | Diurnal cycle | Vertical | Y | Top | Bottom |
| Montipora Incubation 1 | Montipora capricornis | 0.1 | Aquaculture | Diurnal cycle | Vertical | Y | Top | Bottom |
| Montipora Incubation 2 | Montipora capricornis | 0.1 | Aquaculture | Diurnal cycle | Vertical | Y | Top | Bottom |
| Montipora Incubation 2 | Montipora capricornis | 0.1 | Aquaculture | Diurnal cycle | Vertical | Y | Top | Bottom |
| Montipora Incubation 3 | Montipora capricornis | 0.1 | Aquaculture | Diurnal cycle | Horizontal | Y | Side | MiΔΔ le |
| Montipora Incubation 3 | Montipora capricornis | 0.1 | Aquaculture | Diurnal cycle | Horizontal | Y | Side | MiΔΔ le |
| Montipora Incubation 3 | Montipora capricornis | 0.1 | Aquaculture | Diurnal cycle | Horizontal | Y | Side | MiΔΔ le |
| Montipora Incubation 3 | Montipora capricornis | 0.1 | Aquaculture | Diurnal cycle | Horizontal | Y | Side | MiΔΔ le |
| Montipora Incubation 3 | Montipora capricornis | 0.1 | Aquaculture | Diurnal cycle | Horizontal | Y | Side | MiΔΔ le |
| Montipora Incubation 3 | Montipora capricornis | 0.1 | Aquaculture | Diurnal cycle | Horizontal | Y | Side | MiΔΔ le |
| Montipora Incubation 3 | Montipora capricornis | 0.1 | Aquaculture | Diurnal cycle | Horizontal | Y | Side | MiΔΔ le |
| Montipora Incubation 3 | Montipora capricornis | 0.1 | Aquaculture | Diurnal cycle | Horizontal | Y | Side | MiΔΔ le |
| Montipora Incubation 3 | Montipora capricornis | 0.1 | Aquaculture | Diurnal cycle | Horizontal | Y | Side | MiΔΔ le |
| Montipora Incubation 3 | Montipora capricornis | 0.1 | Aquaculture | Diurnal cycle | Horizontal | Y | Side | MiΔΔ le |
| Montipora Incubation 3 | Montipora capricornis | 0.1 | Aquaculture | Diurnal cycle | Horizontal | Y | Side | MiΔΔ le |
| Montipora Incubation 3 | Montipora capricornis | 0.1 | Aquaculture | Diurnal cycle | Horizontal | Y | Side | MiΔΔ le |
| CuracaoTank114 | Rubble | 0.1 | Field collection, 12 m | Darkness | Vertical | N | Top | Bottom |
| CuracaoTank214 | Rubble | 0.1 | Field collection, 12 m | Darkness | Vertical | N | Top | Bottom |
| CuracaoTank315 | Rubble | 0.5 | Field collection, 9 m | Darkness | Vertical | N | Top | Bottom |
| CuracaoTank414 | Rubble | 0.5 | Field collection, 12 m | Darkness | Vertical | N | Top | Bottom |
| CuracaoTank415 | Rubble | 0.5 | Field collection, 9 m | Darkness | Vertical | N | Top | Bottom |
| CuracaoTank512 | Rubble | 0.67 | Field collection, 10 m | Darkness | Vertical | N | Top | Bottom |
| CuracaoTank111 | Sediment | 0.1 | Field collection, 12 m | Darkness | Vertical | N | Top | Bottom |
| CuracaoTank16 | Sediment | 0.33 | Field collection, 1 m | Darkness | Vertical | N | Top | Bottom |
| CuracaoTank211 | Sediment | 0.25 | Field collection, 12 m | Darkness | Vertical | N | Top | Bottom |
| CuracaoTank311 | Sediment | 0.1 | Field collection, 5 m | Darkness | Vertical | N | Top | Bottom |
| CuracaoTank411 | Sediment | 0.25 | Field collection, 5 m | Darkness | Vertical | N | Top | Bottom |
| CuracaoTank110 | Turf | 0.06 | Field collection, 6 m | Darkness | Vertical | N | Top | Bottom |
| CuracaoTank112 | Turf | 0.1 | Field collection, 12 m | Darkness | Vertical | N | Top | Bottom |
| CuracaoTank113 | Turf | 0.1 | Field collection, 5 m | Darkness | Vertical | N | Top | Bottom |
| CuracaoTank115 | Turf | 0.1 | Field collection, 9 m | Darkness | Vertical | N | Top | Bottom |
| CuracaoTank18 | Turf | 0.67 | Field collection, 5 m | Darkness | Vertical | N | Top | Bottom |
| CuracaoTank210 | Turf | 0.06 | Field collection, 6 m | Darkness | Vertical | N | Top | Bottom |
| CuracaoTank213 | Turf | 0.1 | Field collection, 5 m | Darkness | Vertical | N | Top | Bottom |
| CuracaoTank215 | Turf | 0.1 | Field collection, 9 m | Darkness | Vertical | N | Top | Bottom |
| CuracaoTank310 | Turf | 0.06 | Field collection, 6 m | Darkness | Vertical | N | Top | Bottom |
| CuracaoTank312 | Turf | 0.1 | Field collection, 5 m | Darkness | Vertical | N | Top | Bottom |
| CuracaoTank313 | Turf | 0.1 | Field collection, 5 m | Darkness | Vertical | N | Top | Bottom |
| CuracaoTank410 | Turf | 0.06 | Field collection, 6 m | Darkness | Vertical | N | Top | Bottom |
| CuracaoTank412 | Turf | 0.5 | Field collection, 5 m | Darkness | Vertical | N | Top | Bottom |
| CuracaoTank48 | Turf | 0.67 | Field collection, 5 m | Darkness | Vertical | N | Top | Bottom |
| CuracaoTank511 | Turf | 0.5 | Field collection, 9 m | Darkness | Vertical | N | Top | Bottom |
| CuracaoTank515 | Turf | 0.5 | Field collection, 9 m | Darkness | Vertical | N | Top | Bottom |
